# Mitochondrial Respiratory Chain Protein Co-Regulation in the Human Brain

**DOI:** 10.1101/2021.07.19.452923

**Authors:** Caroline Trumpff, Edward Owusu-Ansah, Hans-Ulrich Klein, Annie Lee, Vladislav Petyuk, Thomas S. Wingo, Aliza P. Wingo, Madhav Thambisetty, Luigi Ferrucci, Nicholas T. Seyfried, David A. Bennett, Philip L. De Jager, Martin Picard

**Affiliations:** Department of Psychiatry, Division of Behavioral Medicine, Columbia University Irving Medical Center, New York, USA; Department of Physiology and Cellular Biophysics, Columbia University Irving Medical Center, New York, USA; Center for Translational & Computational Neuroimmunology, Department of Neurology, Columbia University Irving Medical Center, New York, USA; Pacific Northwest National Laboratory, Richland, Washington State, USA; Departments of Neurology and Human Genetics, Emory University, Atlanta, GA, USA; Atlanta VA Medical Center, Decatur, GA, USA and Department of Psychiatry, Emory University, Atlanta, GA, USA; Clinical and Translational Neuroscience Section, Laboratory of Behavioral Neuroscience, National Institute on Aging Intramural Research Program, Baltimore, USA; Longitudinal Study Section, National Institute on Aging, Baltimore, USA; Department of Biochemistry, Emory University, Atlanta, USA; Rush Alzheimer’s Disease Center, Rush University Medical Center, Chicago, Illinois, USA; Department of Neurology, H. Houston Merritt Center, Columbia Translational Neuroscience Initiative, Columbia University Irving Medical Center, New York, USA; New York State Psychiatric Institute, New York, USA

**Keywords:** Mitochondrial respiratory chain, post-mortem brain, Alzheimer disease, proteomics, ROSMAP, BLSA, Banner

## Abstract

Mitochondrial respiratory chain (RC) function requires the stoichiometric interaction among dozens of proteins but their co-regulation has not been defined in the human brain. Here, using quantitative proteomics across three independent cohorts we systematically characterized the co-regulation patterns of mitochondrial RC proteins in the human dorsolateral prefrontal cortex (DLPFC). Whereas the abundance of RC protein subunits that physically assemble into stable complexes were correlated, indicating their co-regulation, RC assembly factors exhibited modest co-regulation. Within complex I, nuclear DNA-encoded subunits exhibited >2.5-times higher co-regulation than mitochondrial (mt)DNA-encoded subunits. Moreover, mtDNA copy number was unrelated to mtDNA-encoded subunits abundance, suggesting that mtDNA content is not limiting. Alzheimer’s disease (AD) brains exhibited reduced abundance of complex I RC subunits, an effect largely driven by a 2-4% overall lower mitochondrial protein content. These findings provide foundational knowledge to identify molecular mechanisms contributing to age- and disease-related erosion of mitochondrial function in the human brain.

## Introduction

The brain contains energivorous networks of interacting neural and glial cells whose energy demands are largely sustained by mitochondria. Mitochondria contain approximately 1,300 proteins (Rath et al., 2020), many of which physically interact with others to produce functional organelles. Key molecular operations in mitochondria are performed by multi-subunit protein complexes, and there is mounting evidence that the stoichiometry of specific protein subunits can influence metabolism and cellular functions. One key example is the mitochondrial respiratory chain (RC), also known as the electron transport chain, which converts reducing equivalents into an electrochemical gradient (Nicholls, 2013) through a process that depends on the stoichiometry of multiple protein subunits.

Perturbing RC protein subunit stoichiometry has functional consequences on mitochondrial and organ function. For instance, mice overexpressing NDUFAB1 (one of the complex I subunits) in skeletal muscles are refractory to obesity and insulin resistance when fed a high-fat diet (Zhang et al., 2019). In addition, overexpression of NDUFAB1 in the heart enhances mitochondrial bioenergetics and limits reactive oxygen species (ROS) production to protect the heart from ischemia-reperfusion injury (Hou et al., 2019). Other studies have shown that the level of expression of NDUFA13 (a complex I subunit also referred to as GRIM-19) is reduced in hepatocellular carcinoma (HCC) cells relative to wildtype controls; and overexpression of NDUFA13 in HCC cells arrests cell proliferation and stimulates apoptosis (Kong et al., 2014). Therefore, the relative abundance of individual subunits in relation to their molecular binding partners is emerging as a contributor to overall organelle and cellular function.

The RC is located within the inner mitochondrial membrane (IMM) and includes five protein complexes. The RC complexes assemble with the help of assembly factors that only temporarily interact with core and accessory protein subunits that are later released from the assembled complexes (Formosa et al., 2018; Mukherjee and Ghosh, 2020). In neurons and other brain cells, RC function ultimately powers not only adenosine triphosphate (ATP) synthesis but also calcium uptake and other biological functions essential to brain function and cognition (Picard and McEwen, 2014; Rangaraju et al., 2019). The ability of the RC to produce adequate amounts of ATP therefore depends on the coordinated expression and stability of its components, the RC protein subunits that are encoded both by the nucleus and by the mitochondrial DNA (mtDNA).

The entry point for the electron transfer is complex I which is the largest RC complex. Complex I contains 45 subunits in humans and assembles in 3 major modular intermediates (Formosa et al., 2018; Garcia et al., 2017). Assembled complex I can further assemble as multimeric supercomplexes with complexes III and IV to enhance electron transfer efficiency (Acín-Pérez et al., 2008; Lapuente-Brun et al., 2013; Letts and Sazanov, 2017; Milenkovic et al., 2017; Vartak et al., 2013). In complex I, 14 of the 45 subunits that form the catalytic centers are referred to as the core or central subunits. Although mitochondrial protein synthesis is considered a highly regulated process (Kummer and Ban, 2021), the extent to which mtDNA-encoded proteins are coregulated to achieve equimolar abundance and the extent to which their abundance is matched to nuclear DNA (nDNA) encoded subunits have not been defined in the human brain.

Mitochondrial RC dysfunction has emerged as a putative cause of neurodegeneration in Alzheimer’s disease (AD) (Holper et al., 2019; Swerdlow et al., 2014) and cognitive decline (Mostafavi et al., 2018; Wingo et al., 2019), as well as Parkinson’s disease (Grünewald et al., 2016), highlighting the importance of understanding brain mitochondrial biology. In particular, we still lack a molecular-resolution understanding of the mitochondrial alterations or recalibrations that may underlie brain dysfunction and AD. Here we investigate mitochondrial RC components in molecularly-profiled post-mortem human brains in relation to AD status and sex. Using proteomic datasets in the human dorsolateral prefrontal cortex (DLPFC), an associative brain region involved in AD, we specifically investigate the co-regulation patterns among known RC protein subunits.

## Results

We characterized the co-regulation of the DLPFC mitochondrial RC proteins by repurposing high-throughput quantitative proteomics measured by liquid chromatography coupled with tandem mass spectrometry (LC-MS/MS). Data from three independent cohorts were analyzed: ROSMAP (N=400) (Bennett et al., 2018; Robins et al., 2019), BLSA (N=47) (Ferrucci, 2008; Swarup et al., 2020; Wingo et al., 2019), and Banner (N=201) (Beach et al., 2015; Swarup et al., 2020; Wingo et al., 2019). Each cohort included both women and men who had either normal cognition, mild cognitive impairment, or AD at the time of death (see **fig. 1A, fig. S1** and **table S1**).

**Fig. 1.**
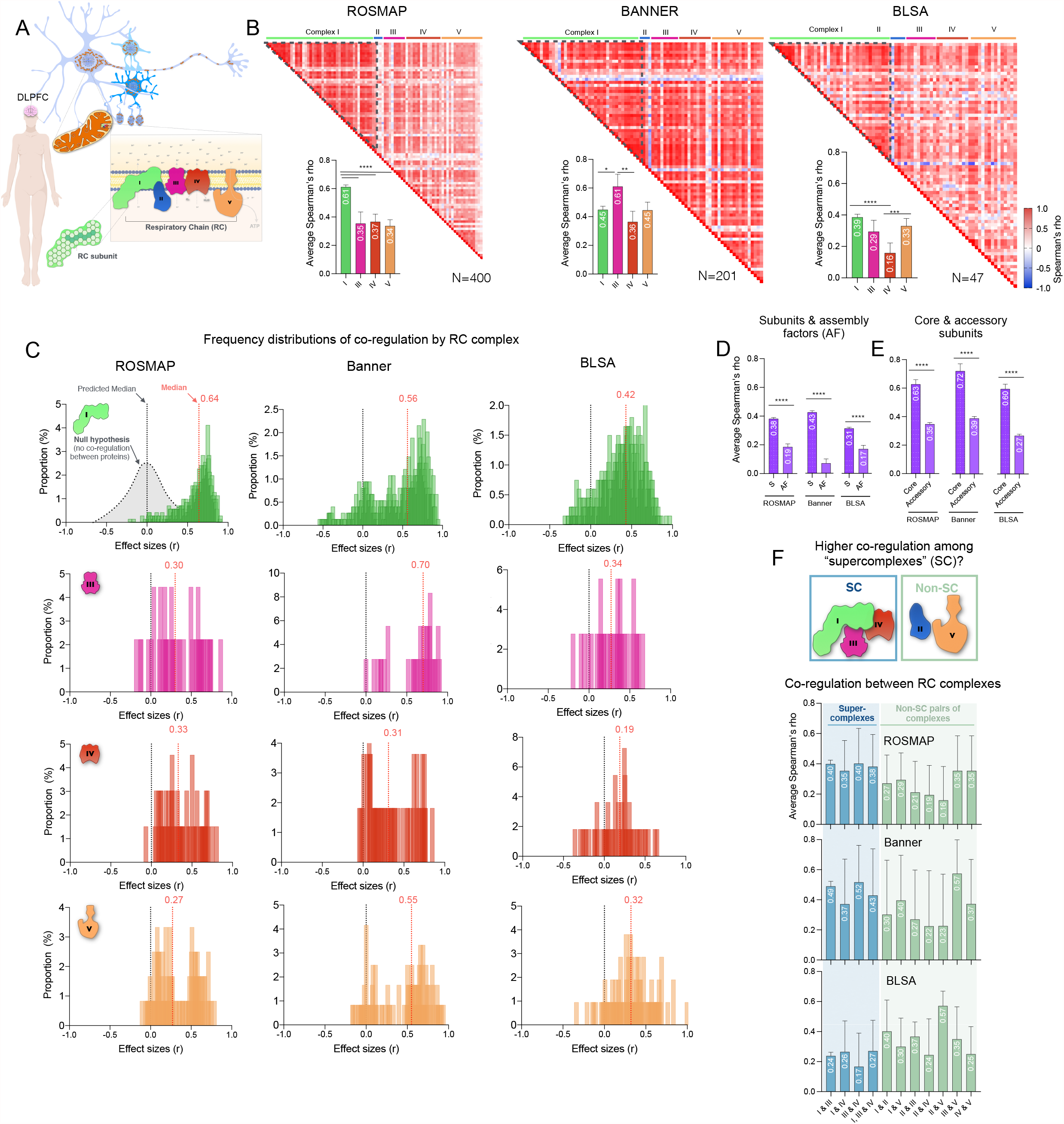
Mitochondrial respiratory chain (RC) complexes proteins co-regulation. (**A**) Study design: we assessed co-regulation of mitochondrial RC proteins (or subunits) in the human dorsolateral prefrontal cortex (DLPFC) across three independent cohorts: ROSMAP (N=400), Banner (N=201) and BLSA (N=47) (for more details see Fig. S1). (**B**) Heatmap of the association between RC nuclear DNA (nDNA) encoded subunits assessed with Spearman rank correlation in ROSMAP, Banner and BLSA. Heatmaps with individual pairs of correlations are presented in Fig S3-S5 and correlation matrices are presented in table S2. Average Spearman’s rho (95% CI) for nDNA encoded subunits for RC complexes I, III, IV, V. (**C**) Distribution of correlation results (Spearman’s rho) for RC complexes I, III, IV, V. Co-regulation of complex II was not studied individually due to its small number of subunits. (**D**) Average Spearman’s rho (95% CI) for nDNA encoded subunits and assembly factors and for (**E**) core and accessory subunits for all RC complexes. (**F**) Average Spearman’s rho (95% CI) for RC complexes part of the hypothesized supercomplexes (SC, I, III, IV) and other possible pairs of RC complexes. P-values from Dunn’s multiple comparisons test are shown in Fig S11A-C. P-values from Mann-Whitney test (D-E) or Kruskal-Wallis test with Dunn’s multiple comparisons test (B), p<0.05*, p<0.01**, p<0.001***, p<0.0001****.

The coverage of RC proteins by all three datasets ranged from 72% to 91% and was largest (91%) in ROSMAP (illustrated in **fig. S1B**). In the ROSMAP dataset, both nDNA-encoded and mtDNA-encoded protein subunits were available, while only nDNA-encoded protein subunits were detected in Banner and BLSA, which used a different proteomics approach to determine protein abundance characterized by less depth than the one used in ROSMAP. Thus, we focused most of our analyses on ROSMAP, the most comprehensively profiled cohort.

Our goal was to assess the degree to which physically-related or non-physically-interacting RC proteins are co-regulated within the human DLPFC. To estimate co-regulation, we performed protein covariation analysis (Kustatscher et al., 2019) by computing the Spearman correlations for each protein pairs and averaging the correlation coefficients (for visualization see **fig. S2**). Additionally, we performed statistical analysis to compare the average co-regulation by subsets of subunits.

### RC protein subunits show co-regulation across RC complexes

Across the three cohorts, the nDNA encoded RC protein subunits exhibited significant co-regulation. This means that brains with higher abundance of a specific RC protein also have more of other related RC proteins, as expected from their assembly in the same complexes (**fig. 1B**, see **fig. S3-S5** for the heatmaps of RC subunits and assembly factors). Computing average co-regulation indices for each RC complex in ROSMAP showed that subunits of Complex I (average across three cohorts r=0.53) were most strongly correlated with each other.

Patterns of co-regulation within each RC complex can also be visualized as frequency histograms indicating the number of protein pairs exhibiting increasing strengths of correlations with other subunits (**fig. 1C**). If co-regulation was absent, we would expect a median distribution value of 0. Here the distribution ranged between r=0.27 - 0.64 which suggests moderate to large co-regulation among RC protein subunits. Between RC complexes, the highest degree of co-regulation was found for Complex I across the three cohorts (average r=0.48). To validate this observation, we also used selected reaction monitoring (SRM), a targeted proteomics approach on a subset of RC proteins in a larger sample of DLPFC samples from the ROSMAP cohort (N=1,228). Again, Complex I subunits showed the highest degree of correlation, suggesting particularly high stability of Complex I and of its subunits in the human brain (**fig. S6**). Of all complexes, Complex IV (average r=0.30) showed the lowest degree of co-regulation. Complex II was not assessed individually since it includes too few (four) subunits.

Across the RC proteins, assembly factors are not part of the final assembled RC complexes. If the physical interactions of subunits among assembled complexes contribute to co-regulation, assembly factors should exhibit lower co-regulation. Indeed, compared to RC subunits across all complexes (average r=0.37), we found approximately two-third lower co-regulation for assembly factors (average r=0.14) (**fig. 1D**). Among actual RC protein subunits, *accessory* subunits, which represent the structural components of the RC, showed lower co-regulation (average r=0.33) than *core* subunits responsible for performing the catalytic activities (average r=0.65) (**fig. 1E**).

### Co-expression of RC subunits in RNAseq

To examine if the observed co-regulation of RC subunits is a feature only present at the protein level, or if transcriptional activity (i.e., gene expression) could contribute to these effects, we also examined the co-expression of RC components based on mRNA transcripts. Bulk RNAseq data was examined in the DLPFC (N=1,092), as well as two other brain regions: the posterior cingulate cortex (PCC, N=661) and the anterior caudate (AC, N=731).

Similar to our observations at the protein level, in the DLPFC, PCC and AC, individuals with higher mRNA levels for specific subunits tended to also have higher levels of other related transcripts, consistent with the co-expression of RC genes as part of a common genetic program (**fig. S7-9**). In the DLPFC, co-regulation estimates based on proteins were slightly weaker (−7.3%) than RNA levels (**fig. S7C**). Thus, these data suggest that co-regulation of RC subunits occurs at both the protein and transcript levels.

### Partial evidence for the existence of RC supercomplexes in the human brain

Research in drosophila and mammalian model systems suggests that the RC complexes I, III and IV assemble as supercomplexes (Acín-Pérez et al., 2008; Lapuente-Brun et al., 2013; Letts and Sazanov, 2017; Milenkovic et al., 2017; Vartak et al., 2013). If supercomplexes (SC) also existed in the human brain, we would expect RC subunits between these complexes to be more highly co-regulated than with subunits of non-SC subunits (complex II and V). We therefore asked if pairs of subunits derived from complex I-III-IV are more strongly co-regulated with each other than other RC complexes (II, V). In the more comprehensive ROSMAP dataset, we find that co-regulation between SC RC complexes was 46% higher compared to non-SC RC complexes in ROSMAP (**fig. 1E, fig. S11A-C**). Similarly, in the Banner dataset, the co-regulation between complex I, III and IV was 43% higher (p<0.01) than for complex I and II. This effect, however, was not observed in the BLSA cohort, perhaps due to lower power in detecting a difference in co-regulation or possibly due to other inherent differences in participants characteristics between cohorts. Higher co-regulation of SC was also observed at the RNA levels across three brain regions, DLPFC, PCC, AC (ROSMAP) (**fig. S10, fig. S11D-F**).

### Variation in RC protein abundance, mitochondrial mass, and mtDNA density

To understand what drives the variation in RC subunits in the DLPFC, we assessed the correlation between nDNA- and mtDNA-encoded RC subunits and i) markers of mitochondrial content, ii) mtDNA copy number (mtDNAcn), and iii) neuronal proportion (**fig. 2 A,B**, see table S1 for the descriptive statistics). We built an index of mitochondrial content using the average of the scaled values of four housekeeping proteins including the translocator of the outer membrane (TOMM20.Q15388), voltage-dependent anion channel 1 (VDAC1.P21796, also known as porin), and citrate synthase (CS.O75390, CS.H0YIC4). As expected, brains with higher general mitochondrial content also had higher RC protein subunits abundance, as indicated by the positive correlation of the mitochondrial content index and individual proteins (**fig. 2 A,B**). Based on these estimates, overall mitochondrial content explains ∼23% of the variance in RC protein abundance. The unexplained variance suggests that mitochondria may vary in their RC protein density (i.e., how much RC each mitochondrial unit contains) (McLaughlin et al., 2020).

**Fig. 2.**
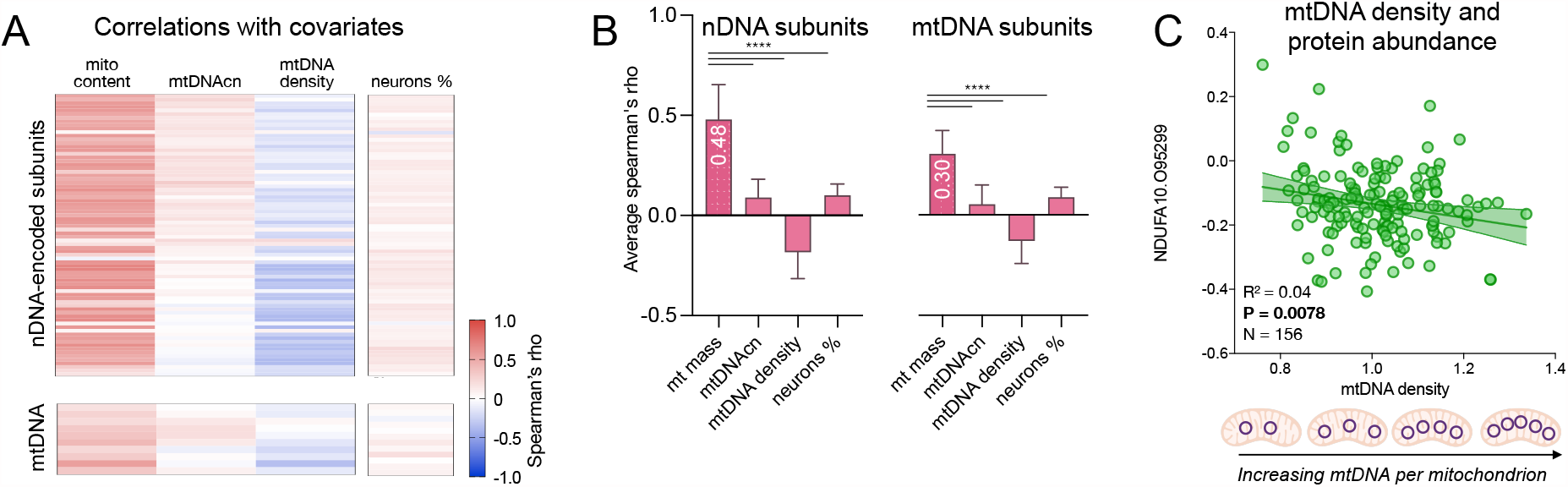
Relationship between mitochondrial respiratory chain (RC) protein abundance, mitochondrial mass, mtDNA density, mitochondrial DNA copy number (mtDNAcn) and neuron proportion. Association between RC nuclear DNA (nDNA) and mtDNA encoded subunits abundance and mitochondrial content (average of the scaled values of CS.O75390, CS.H0YIC4, TOMM20.Q15388, VDAC1.P21796), mtDNAcn, mtDNA density (mtDNAcn/mito content) and neuron proportion (neuron %) shown as (**A**) heatmaps of Spearman’s rho (**B**) and as average Spearman’s rho (95% CI). P-value from Kruskal-Wallis test with Dunn’s multiple comparisons test (B, left) or one-way ANOVA and Tukey’s multiple comparison test (B, right), p<0.0001****. (**C**) Example of scatterplot of the association between a complex 1 subunit (NDUFA10.O95299) and mtDNA density. Results from simple linear regression. ROSMAP, N=124-400

Modest positive associations were found between mtDNAcn quantified from whole genome sequencing (WGS) (N=454) and either nDNA- and mtDNA-encoded RC subunits, such that mtDNAcn accounted for only 0.1 % of the variance in RC protein abundance. Thus, mtDNAcn alone or in combination with mitochondrial content, accounts for at most 24% of inter-individual differences in RC protein abundance in the DLPFC (**fig. 2**).

To examine whether the number of mtDNA copies per mitochondrion is a fixed quantity or varies between individuals, we divided mtDNA copies by the mitochondrial content index. The resulting metric of mtDNA density per mitochondrion differed by up to 6% between individuals with the highest and lowest mtDNA density, establishing the range and natural variability of this parameter in the human DLPFC. Interestingly, brains with greater mtDNA density tended to have lower total RC protein abundance, including mtDNA-encoded proteins (**fig. 2 A,B** and **C** for an example). This suggests that increased mtDNA copies for a given mitochondrial mass does not lead to higher mitochondrial RC abundance. Rather, elevated mtDNA per mitochondrion might be the result of compensatory mechanisms activated by low RC protein concentration, as seen in genetic cases of mitochondrial dysfunction (Giordano et al., 2014; Yu-Wai-Man et al., 2010).

Neurons are rich in mitochondria, and differences in the number of neurons per unit of brain tissue that may occur secondary to neuronal loss with neurodegeneration (Gómez-Isla et al., 1997) could theoretically contribute to RC chain abundance. As an exploratory analysis, we also examined to what extent RC protein abundance may be driven by neuronal abundance. In other words: do brains with more neurons relative to other cell types have higher RC protein abundance? On average less than 5% of the variance in RC protein abundance was explained by neuron density derived from RNA-seq within the DLPFC (**fig. 2 A,B**), ruling out neuronal density as a major confounding factor in these analyses, and confirming that non-neuronal cells make a significant contribution to the brain proteomics signal.

### Evidence of mito-nuclear crosstalk in human DLPFC

Both nDNA- and mtDNA-encoded protein subunits physically interact in the RC, and coordination of mitochondrial and nuclear genomes (i.e., mitonuclear crosstalk) is considered an important determinant of mitochondrial biology (Kim et al., 2018; Quirós et al., 2016). To indirectly quantify evidence of co-regulation within and across genomes, we assessed the co-regulation among nDNA-encoded subunits, and separately among mtDNA encoded subunits in ROSMAP. Moreover, to estimate to what extent mito-nuclear crosstalk produces coordinated levels of proteins encoded in both genomes, we specifically quantified the co-regulation *between* nDNA- and mtDNA-encoded subunits (**fig. 3A**).

**Fig. 3.**
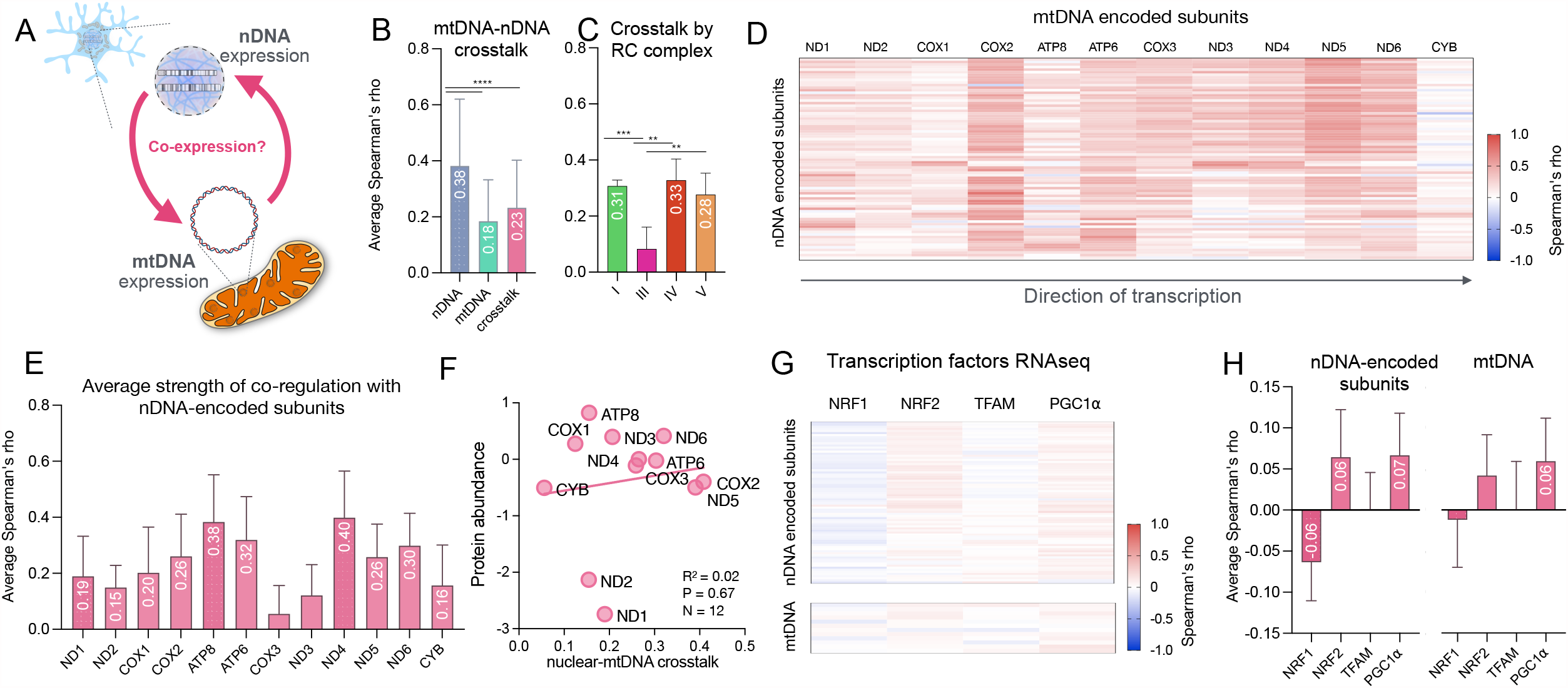
mtDNA and nuclear DNA (nDNA) encoded mitochondrial respiratory chain (RC) proteins co-regulation, nuclear-mtDNA crosstalk and transcription factors. (**A**) Diagram of nuclear-mtDNA crosstalk. (**B**) Average Spearman’s rho (95% CI) between nDNA encoded subunits, between mtDNA encoded subunits and for the crosstalk between nDNA and mtDNA subunits. (**C**) Average Spearman’s rho of nuclear-mtDNA crosstalk for RC complexes I, III, IV, V. (**D**) Heatmap and (**E**) average Spearman’s rho (95% CI) of nuclear-mtDNA crosstalk presented by mtDNA encoded subunits’ order of transcription. Note that ND4L is not shown (data not available). P-values from Dunn’s multiple comparisons test are shown in Fig. S12, (**F**) Relationship between mtDNA encoded RC protein abundance and nuclear-mtDNA crosstalk. Results from simple linear regression. (**G**) Heatmap and (**H**) average Spearman’s rho (95% CI) of RC subunits protein abundance and transcription factor gene expression (NRF2 corresponds to GABPB1 gene). P-values from Dunn’s multiple comparisons test are shown in Fig S13. ROSMAP, N =124-400. P-value from Kruskal-Wallis test with Dunn’s multiple comparisons test, p<0.05*, p<0.01**, p<0.001***, p<0.0001****.

The extent of co-regulation among nDNA-encoded subunits was more than double (r=0.38) that of mtDNA subunits (r=0.18) (**fig. 3B**). The degree of crosstalk between nDNA- and mtDNA-encoded subunits was also low (r=0.23) (**fig. 3B**). Among individual RC complexes, the lowest degree of nDNA-mtDNA crosstalk was found for complex III (r=0.08, <1% of shared variance) and the highest was found for complex I and IV (r=0.31-0.33, ∼10% shared variance) (**fig. 3C**). For mtDNA-encoded genes, the observed strength of co-regulation was not associated with the order of the genes or their distance from the initiation of transcription sites. Co-regulation with nDNA-encoded subunits was largest for ND1, COX2, and ND5, and lowest for CYTB (**fig. 3 D,E, fig. S12**). Co-regulation differences between proteins were not confounded by their abundance (**fig. 3F**), although we note the existence of a possible trend (excluding ND1 and ND2) whereby lower abundance proteins may exhibit higher mito-nuclear crosstalk.

We then explored how much the observed inter-individual differences in the abundance of nDNA- and mtDNA-encoded subunits are explained by canonical transcription factors (TFs) believed to mediate mito-nuclear crosstalk. We examined TFs acting either on the nuclear (NRF1, NRF2, PGC1a) or mitochondrial (TFAM) genomes, and quantified their expression measured by RNA-seq with RC protein and transcript abundance. TFs expression did not correlate with RC subunits abundance (all rs<0.1) (**fig. 3G,H fig. S13**) but expression of NRF2 and TFAM correlated moderately with RC transcript abundance (**fig. S14**). Importantly, we also found either no or weak associations between RC transcript (RNA-seq) (Raj et al., 2018) and protein abundance levels (**fig. S15**).

### Complex I proteins co-regulation by topology

Since co-regulation was highest for complex I in ROSMAP as well as in BLSA, and since it is the largest RC complex with high-fidelity three-dimensional maps of its subunits, we refined our covariance analysis of complex I subunits according to their topological location (**fig. 4A**). This analysis focused on the ROSMAP cohort because it is the largest, has the best global coverage, and included both nDNA and mtDNA-encoded proteins. Moreover, ROSMAP contains a substantial number of individuals with either normal cognition or with an AD diagnosis, allowing to examine associations with AD.

**Fig. 4.**
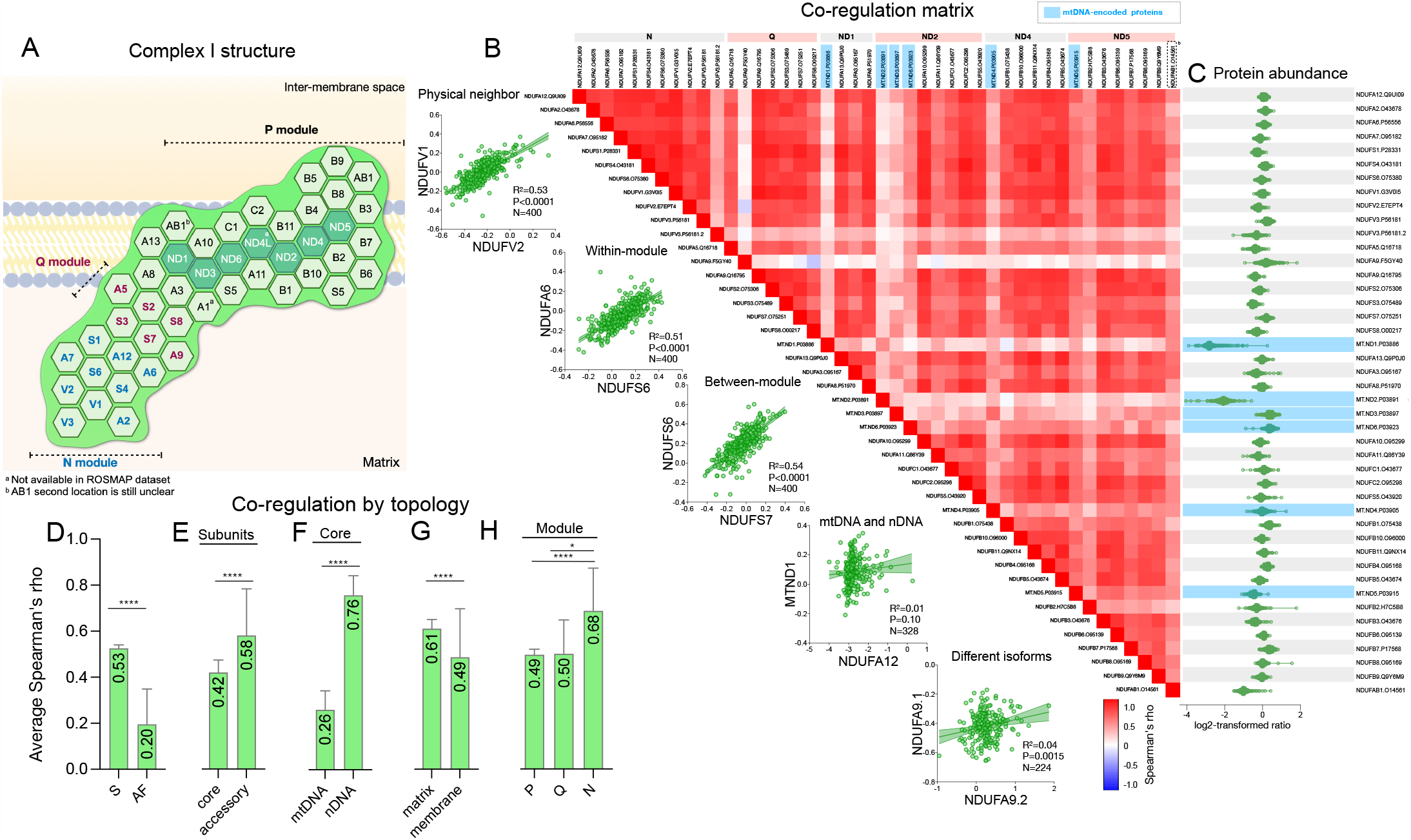
Mitochondrial respiratory chain (RC) complex I proteins co-regulation. (**A**) Diagram of complex I protein subunits topological location. (**B**) Heatmap of complex I protein co-regulation arranged by modules and example of individual association (results from simple linear regression). (**C**) Complex I subunit abundance (raw values). Average Spearman’s rho (95% CI) coefficient for complex I (**D**) subunits and assembly factors, (**E**) core and accessory subunits (**F**) nuclear DNA (nDNA) or mtDNA encoded subunits (**F**) subunits by localization and (**G**) by module. ROSMAP, N=124-400.P-value from unpaired t-test, Mann-Whitney test, or Kruskal-Wallis test with Dunn’s multiple comparisons test, p<0.05*, p<0.01**, p<0.001***, p<0.0001****.

**Fig. 4B** displays the detailed correlation matrix of RC complex I subunits arranged by their topological location. **Fig. 4C** shows the corresponding protein abundance for each subunit, illustrating that co-regulation is not driven by protein content (**fig. 4B,C**). As observed across all RC complexes, structural subunits that physically assemble into the protein complex showed substantially higher co-regulation than assembly factors (**fig. 4D**). Among RC subunits, accessory subunits had 30% larger co-regulation than core subunits, but this difference is mainly driven by mtDNA-encoded subunits that show substantially lower co-regulation than nDNA encoded subunits (r=0.26 vs r=0.76).

Next, we assessed whether RC protein co-regulation varies according complex I subunit topological location. Mitochondrial complex I is an ‘L’ shaped enzyme composed of a hydrophilic arm that protrudes into the matrix, and a hydrophobic arm that is embedded within the inner membrane. Matrix-located subunits showed 24% higher co-regulation than membrane subunits (**fig. 4G**). Proteins located in the N module (r=0.68) showed greater co-regulation than proteins located in the Q (r=0.50) or P module (r=0.49) (**fig. 4H**). These topologically-sensitive covariance patterns in the human DLPFC provide information about the relative co-regulation and/or stability, or regulated turnover of all proteins within each module *in vivo* in the human brain.

We also note that some nDNA-encoded complex I subunits exhibited particularly low co-regulation with any of the other subunits, such as NDUFA9.F5GY40, and NDUFV3.P56181.2. In both cases, these proteins have a related isoform that has high co-regulation with neighboring and other subunits, suggesting that these isoforms are generally not incorporated in the assembled complex I in the human DLPFC.

### Complex I proteins co-regulation by sex

Sex differences in energy metabolism have been observed in humans (Mittelstrass et al., 2011), and studies of various aspects of mitochondrial biology in model organisms suggest that mitochondria in males exhibit lower mitochondrial content and ATP production capacity compared to females (Ventura-Clapier et al., 2017). Therefore, we examined whether males and females exhibit differences in complex I co-regulation. To explore this question, we computed the same covariance matrix for all complex I subunits as in figure 4, but separately for men (**fig. S16A)** and women (**fig. S16B)**, then generated a difference matrix (see **fig. S16C** for the matrix of women-men difference where correlations that differ between the sexes appear in darker colors). Differences in co-regulation ranged from -0.43 to 0.36 (average=0.02). We then ran an interaction analysis (correlations of protein-protein *x* sex) on the top 100 most different correlation pairs, of which the top-15 most significant are shown in **fig. S16D**. Although we did not find any systematic difference between women and men, these data suggest potential differences among specific protein pairs that could be of interest to future studies of sex differences. For example, ND2 and NDUFA13 (both P module subunits) were strongly correlated in women, but not in men.

### Complex I proteins co-regulation by clinical diagnosis

In relation to AD, multiple studies have found that Complex I enzymatic activity is reduced in AD (see (Holper et al., 2019) for a meta-analysis). Therefore, we hypothesized that this enzymatic deficiency could be associated with altered complex I protein co-regulation. To explore differences in complex I protein co-regulation according to clinical diagnosis of AD in ROSMAP, we assessed separately the correlation between subunits in individuals with normal cognitive function at death (NCI, N=170, **fig. S16F**) and in individuals with AD (N=123, **fig. S16E)**; then, we assessed the difference in effect size between NCI and AD (**fig. S16G**). AD was defined using the NIA-Reagan criteria for the pathologic diagnosis of AD (high or intermediate likelihood of AD) (Bennett et al., 2006) and final clinical diagnosis of cognitive status (AD and no other cause of cognitive impairment) as described previously (Klein et al., 2020). Differences in protein-protein correlations ranged from -0.32 to 0.30 (average=-0.02). Then, we ran an interaction analysis on the top 100 most different correlation pairs, of which the top-15 most significant are shown in **fig. S16H**. As for sex differences, there was no systematic difference in complex I co-regulation between AD and NCI, indicating that co-regulation of complex I subunits is unlikely to underlie enzymatic activity defects in AD.

### Complex I proteins abundance by sex

In the absence of systematic co-regulation differences between women and men, or between AD and NCI, we reasoned that there may be a coordinated up or down regulation in protein abundance between women/men and AD/NCI. Overall, men and women did not show a difference in the overall complex I protein abundance. The average standardized effect size Hedge’s g was 0.004, indicating the absence of a difference in complex I subunit abundance in relation to sex (**fig. S17A)**.

### Complex I proteins abundance by clinical diagnosis

In contrast, the analysis by AD status revealed a general downregulation of several complex I subunits. Of the 44 complex I subunits available, 95% were significantly lower in AD than in NCI (p<0.0001, Chi-square compared to chance (50%)), and none were upregulated. The protein with the largest effect size was NDUFB7, decreased by an average of 7%, representing a medium to large effect size (g=0.70) (**fig. S17B**).

### RC proteins abundance and mitochondrial mass by sex and clinical diagnosis

We next determined whether the observed AD/NCI difference were i) specific to complex I or whether they generalized to other RC complexes, ii) observed in other cohorts, and ii) driven by reduced mitochondrial mass in AD brains. Therefore, we studied men/women and AD/NCI differences in the five RC complexes (**fig. 5**) using the average fold change in individual subunits **(fig. 5A, D**). We also used summary scores representing mitochondrial content and average RC abundance by complex, adjusted or not for mitochondrial content **(fig. 5B, E**). While we did not observe any systematic difference in RC abundance between women and men, mitochondrial content tended to be 0.3-1.6% higher in men than in women **(fig. 5C**). Relative to NCI, AD brains had on average 2.6-3.7% lower abundance of RC complexes, but this effect was largely driven by a 1.9-3.9% lower mitochondrial content across all three cohorts. Therefore, when accounting for mitochondrial content, mitochondria from AD and NCI do not appear to differ in their RC protein abundance **(fig. 5F**). However, at the brain tissue level, our proteomics results are consistent with and could explain the reported loss of cortical mitochondrial oxidative capacity in AD (Holper et al., 2019).

**Fig. 5.**
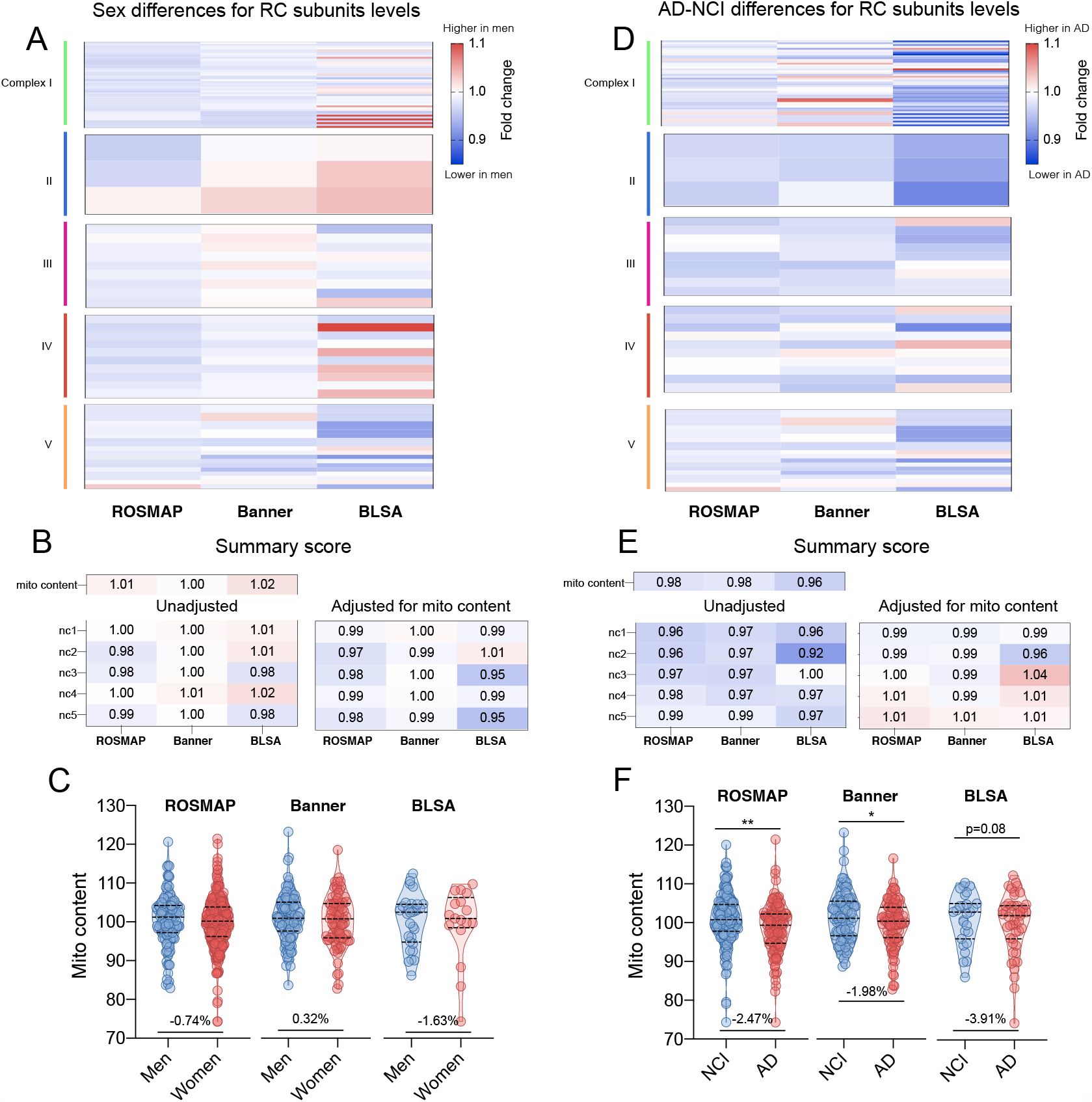
Difference in mitochondrial respiratory chain (RC) proteins abundance according to sex and clinical diagnosis. Difference between men and women in proteins abundance shown as fold change for the (**A**) RC individual subunits and (**B**) mitochondrial content and RC summary scores. (**C**) violin plot of mito content by sex. P-value from one-way ANCOVA adjusted for Alzheimer’s disease (AD) status and age at death. Difference between individuals with no cognitive impairment (NCI) and AD at death in RC proteins abundance shown as (**D**) RC individual subunits, (**E**) mitochondrial content and RC summary scores. (**F**) violin plot of mito content by AD status. P-value from one-way ANCOVA adjusted for sex and age at death (as well as years of education for ROSMAP and BLSA). Scores were adjusted for post-mortem interval, age at death, study, and batch. *p<0.05, **p<0.01. Detailed results are shown in supplementary table S4.

## Discussion

We deployed untargeted proteomics and covariance analysis to identify patterns of co-regulation among mitochondrial RC proteins in the human DLPFC. A key aspect of this study was the integration of known biological and topological information, particularly for complex I, to inform the interpretation of protein co-regulation. Consistent with the physical assembly of *bona fide* subunits into physical complexes that have coordinated turnover (Szczepanowska et al., 2020), relative to assembly factors that assist in the assembly but are not part of the final assembled complexes, our approach reliably detected higher co-regulation of assembled subunits than assembly factors across all examined cohorts. Focusing on the largest cohort, ROSMAP, which also has the highest proteomic coverage across both mtDNA-and nDNA-encoded subunits, we find that the largest RC complex, complex I, exhibits a high level of co-regulation relative to other complexes. Our data also identifies interesting patterns of regulation, notably the lack of association between RC subunits and mtDNA abundance in brain tissue.

Our data aligns well with a recent model of complex I maintenance in cultured cells (Szczepanowska and Trifunovic, 2021) and extends its potential applicability to the human brain. In particular, our analysis of protein-protein co-regulation and abundance for different complex I subunits and isoforms in 400 individuals provides a platform to further examine mitochondrial RC regulation. For example, specific isoforms exhibiting relatively normal abundance but no co-regulation with other subunits may not assemble *in vivo*. The high co-regulation of groups of subunits, such as those contained within the N-module of complex I, could be attributable to its high turnover as a single module (Szczepanowska et al., 2020). The conservation of this pattern from cultured cells to the human brain suggests that this process is robust and possibly a general principle of mitochondrial RC maintenance.

Another noteworthy observation is the lack of apparent co-regulation between RC subunits encoded in the nuclear and mitochondrial genomes. Although in theory transcription and translation of gene products in both genomes are coupled via several mito-nuclear communication mechanisms (Quirós et al., 2016), our results indicate a disconnect (i.e., lack of effective co-regulation) between RC proteins, which may point to protein regulation mechanisms operating relatively independently between mitochondrial and nuclear compartments in the human DLPFC. In other words, brain mtDNA-related protein synthesis could occur relatively independently from that of cytoplasmic nDNA-encoded proteins. Whether this potential mito-nuclear protein imbalance in the human brain could contribute to activating compensatory mechanisms, such as the mitochondrial unfolded protein response (UPR^mt^) (Houtkooper et al., 2013), could also be examined in relation to proteotoxic stress and brain pathology. This point also may be consistent with the fact that mtDNAcn was not found to be correlated with mtDNA- or nDNA-encoded RC proteins, possibly implicating translational regulation (Kummer and Ban, 2021) and/or equally influential degradation and turnover processes, as the main drivers of RC protein abundance, rather than the number of mtDNA molecules per cell. Moreover, potential differences in compartment-specific translation (Kummer and Ban, 2021) and degradation (Chung-ha et al., 2014) (e.g., in axons, synaptic terminals, cell bodies) that could not be resolved in our work should also be considered in future studies.

In relation to mtDNA abundance, combining precise estimates of mtDNA copies per cell from WGS and proteomic estimates of mitochondrial abundance showed two main points. First, that mtDNAcn is not a good surrogate for mitochondrial abundance. It is recognized that mtDNAcn is under the influence of a multitude of factors (Filograna et al., 2020) and is generally not a good marker of mitochondrial content, at least in human skeletal muscle (Larsen et al., 2012). Our data corroborate this notion in the human DLPFC. Furthermore, by quantifying the number of mtDNA copies per mitochondria, or mtDNA density, we find that mitochondria with greater mtDNA density have lower RC protein abundance. This inverse association could be similar to what occurs in mitochondrial disease, where poorly functioning mitochondria (few RC proteins) activate signaling cascades to produce a compensatory upregulation of mtDNA copies as an attempt to restore mitochondrial oxidative capacity – an effect observed at the tissue and single-cell level (Giordano et al., 2014; Vincent et al., 2018; Yu-Wai-Man et al., 2010). A meta-analysis showed that aging (the majority of our population) and AD are associated with lower complex I and IV enzymatic activities (Holper et al., 2019), suggesting that functional impairments could indeed exist in a portion of the brains examined here. Combined with the marginally lower mitochondrial content detected in our data, this could possibly drive elevation in mtDNA density in the DLPFC. Therefore, we speculate that combined with other measures of mitochondrial function (such as a ratio reflecting mtDNA abundance per mitochondrial mass), mtDNA density could represent a potential marker of mitochondrial dysfunction to be examined in future studies.

Finally, similar to previous work (de Sousa Abreu et al., 2009; Maier et al., 2009) and to recent observations across human tissues showing that RNA transcript levels are generally poorly correlated with protein abundance (Jiang et al., 2020), we found no evidence of correlation between RNA and protein levels for most RC proteins. Although RNA integrity is generally high in all postmortem samples processed (Mostafavi et al., 2018), we cannot entirely rule out the influence of uncontrolled confounds, such as degradation of RNA transcripts during the post-mortem interval relative to greater protein stability, which would contribute to diminish RNA-protein correlations. Regardless, our analyses of the human brain proteome show conserved and expected co-regulation patterns consistent with RC composition and topology. Overall, this comprehensive analysis of RC subunit co-regulation patterns in a large cohort of women and men, with or without AD, adds molecular-resolution data to further examine the basis for normal and abnormal mitochondrial function in the human brain.

## Supporting information

S1 Descriptives

S2 Correlation matrices

S3 Correlation matrices stratified

S4 GLM

S5 annotated protein list

## Acknowledgments

We acknowledge participants of the ROS and MAP, Banner and BLSA studies. Support for this work was provided by NIH grants P30AG16101 (DAB), R01AG15819 (DAB), R01AG17917 (DAB), U01AG46152 (PLD, DAB), U01AG61356 (PLD, DAB), R01AG036836 (PLD), U01AG061357 (SNT), R01AG061800 (SNT), RF1AG062181 (SNT), GM119793 (MP) and MH122706 (MP), the Intramural Program of the National Institute on Aging (MT), a NARSAD young investigator award (CT) and the Nathaniel Wharton Fund (MP).

## Author contributions

CT, MP and EOA designed the analytic plan. CT performed the data analysis. CT, MP and EOA drafted the original draft the manuscript. All authors participated to the writing, review and editing of the manuscript.

## Competing Interests

The authors declare no competing interests.

## STAR Methods

### RESOURCE AVAILABILITY

| RESOURCE | SOURCE | IDENTIFIER |
| --- | --- | --- |
| **Database** |  |  |
| ROSMAP TMT proteomics | (Robins et al., 2019) | syn17015098 |
| ROSMAP SRM proteomics | (Yu et al., 2018) | syn10468856 |
| ROSMAP RNA-seq | (Raj et al., 2018) | syn3388564 |
| ROSMAP mtDNAcn from WGS | (Klein et al., 2021) | syn25618990 |
| Banner Raw Proteomic Data | (Swarup et al., 2020) | syn7170616 |
| BLSA Raw Proteomic Data | (Seyfried et al., 2017) | syn3606086 |
| **Software and algorithms** |  |  |
| RStudio | https://www.rstudio.com/ | v1.4 |
| R | https://cran.r-project.org/ | v4.0.4 |
| Prism | https://www.graphpad.com/ | V9.1.1 |

#### Lead contact

Further information and requests for resources should be directed to and will be fulfilled by the lead contact, Martin Picard.

#### Materials availability

This study did not generate new unique materials or reagents.

#### Data and code availability

Access to full datasets from the ROS and MAP studies can be requested on the website: https://www.radc.rush.edu/. The molecular data used in this study is available through the synapse.org AMP-AD Knowledge Portal: www.synapse.org

### EXPERIMENTAL MODEL AND SUBJECT DETAILS

We used data from ROSMAP (N=400) (Bennett et al., 2018; Robins et al., 2019; Yu et al., 2020), BLSA (N=47) (Ferrucci, 2008; Swarup et al., 2020; Wingo et al., 2019), and Banner (N=201) (Beach et al., 2015; Swarup et al., 2020; Wingo et al., 2019). These studies have measured post-mortem DLPFC protein abundance using untargeted proteomics. Clinical characteristics of the study participants included in the analysis are shown in supplementary table S1.

### ROSMAP

### Participants

The Rush Memory and Ageing Project (MAP) and the Religious Orders Study (ROS) (A Bennett et al., 2012a; A Bennett et al., 2012b; Bennett et al., 2018) are two ongoing cohort studies of aging and dementia in older persons. The ROS study enrolls Catholic nuns, priests, and brothers, from about 40 groups across the United States. The MAP study enrolls participants primarily from about 40 retirement communities throughout northeastern Illinois, with additional diverse participants via individual home visits. Participants in both cohorts were free of known dementia at study enrollment and agreed to annual evaluations and organ donation on death. Both studies were approved by an Institutional Review Board of Rush University Medical Center and all participants signed an informed consent, Anatomical Gift Act, and a repository consent to share data and biospecimens.

The clinical diagnosis of Alzheimer’s Disease (AD) was based on the analysis of the annual clinical diagnosis of dementia by the study neurologist blinded to post-mortem data. Post-mortem AD pathology was assessed as described previously (Bennett et al., 2006; Bennett et al., 2003), and AD classification was defined based on the National Institutes of Ageing-Reagan criteria (Hyman and Trojanowski, 1997). Dementia status was coded as no cognitive impairment (NCI), mild cognitive impairment (MCI), or Alzheimer’s dementia (AD) from the final clinical diagnosis of dementia. Pathologic AD was defined using the NIA-Reagan criteria for the pathologic diagnosis of AD (high or intermediate likelihood of AD) (Bennett et al., 2006) and final clinical diagnosis of cognitive status (AD and no other cause of cognitive impairment) as described previously (Klein et al., 2020). MCI refers to those persons with cognitive impairment but who did not meet criteria for dementia.

#### Additional cohorts: Banner and BLSA Banner Participants

As a first replication sample, we used proteomic data of post-mortem DLPFC brain tissue from the Banner Sun Health Research Institute’s Brain and Body Donation Program. Data was available from 101 cognitively normal (controls) and 100 Alzheimer’s disease (AD) cases (43% women). Subjects were enrolled as cognitively unimpaired volunteers from the retirement communities (Beach et al., 2015) or as patients by neurologists. The Banner study was approved by the Banner Sun Health Research Institute Institutional Review Board-approved. Participants in the Banner dataset provided informed consent for clinical assessments during life, brain donation after death, and usage of donated biospecimens for approved future research (Beach et al., 2015).

Post-mortem neuropathological evaluation was performed at Banner Sun Health Research Institute with the assessment of amyloid plaque distribution according to CERAD criteria and the neurofibrillary tangle pathology assessed with Braak staging. Control cases were defined as cognitively healthy within on average 9 months of death when they presented low CERAD (0.13 ±0.35) and Braak (2.26 ±0.94) measures of amyloid and tau neuropathology. Participants were classified as AD cases when evaluated as demented at the last clinical research assessment, and when the brains showed high CERAD (2.9 ±0.31) and Braak (5.4 ±0.82) scores (consistent with moderate to severe neuropathological burden).

#### BLSA Participants

As a second replication sample, we used proteomic data performed on DLPFC brain tissue samples from the National Institute on Aging’s Baltimore Longitudinal Study of Aging. Data was available for 13 cognitively healthy individuals, 14 asymptomatic AD and 20 AD cases (36% women). The BLSA study was approved by the Institutional Review Board and the National Institute on Aging. Human research at the National Institutes of Health (NIH) and the BLSA participants provided written informed consent (Ferrucci, 2008).

Post-mortem neuropathological evaluation was performed at the Johns Hopkins Alzheimer’s Disease Research Center with the Uniform Data Set. Assessment included amyloid plaque distribution according to the CERAD criteria and neurofibrillary tangle pathology assessed with the Braak staging. Control cases were defined as cognitively healthy within on average 9 months of death when they presented low CERAD (0.13 ±0.35) and Braak (2.26 ±0.94) measures of amyloid and tau neuropathology. Participants were classified as AD cases when evaluated as demented at the last clinical research assessment, and when the brains showed high CERAD (2.9 ±0.31) and Braak (5.4 ±0.82) scores (consistent with moderate to severe neuropathological burden). Asymptomatic AD cases were cognitively normal proximate to death, and had high CERAD (2.1 ±0.52) and moderate Braak (3.6 ±0.99).

## METHOD DETAILS

### ROSMAP

#### Tandem mass tag (TMT) isobaric labeling mass spectrometry

Untargeted proteomics was measured from post-mortem DLPFC (Broadman area 9) tissue of 400 (70% women) individuals including 168 cognitively normal individuals, 101 MCI, 123 AD and 8 had other cause of dementia, using tandem mass tag (TMT) isobaric labeling mass spectrometry methods for protein identification and quantification as previously described (Robins et al., 2019). A total of 12,691 unique proteins (7,901 after quality control) were detected.

Briefly, tissue homogenization was performed (following the method described in (Ping et al., 2018)) before protein digestion. An equal amount of protein from each sample was aliquoted and digested in parallel to serve as the global pooled internal standard (GIS) in each TMT batch. Samples were randomized by co-variates (age, sex, PMI, diagnosis, etc.) into 50 batches (8 cases per batch) before TMT labeling. The samples (N=400) and the pooled global internal standards (GIS) (N=100) were labelled using the TMT 10-plex kit (ThermoFisher 90406) as previously described in (Johnson et al., 2018; Ping et al., 2018). In each batch, TMT channels 126 and 131 were used to label GIS standards and the 8 middle TMT channels were used for individual samples following randomization. The labelling was followed by high-pH fractionation on an Agilent 1100 HPLC system (as described in (Mertins et al., 2018; Robins et al., 2019)). Peptide eluents were separated on a self-packed C18 (1.9 μm, Dr. Maisch, Germany) fused silica column (25 cm × 75 μM internal diameter (ID); New Objective, Woburn, MA) by a Dionex UltiMate 3000 RSLCnano liquid chromatography system (ThermoFisher Scientific) and monitored on an Orbitrap Fusion mass spectrometer (ThermoFisher Scientific). The RAW files were analyzed using the Proteome Discoverer suite (version 2.3, ThermoFisher Scientific). The protein abundances were log2 transformed and regressed for age at death, postmortem interval, study and batch for downstream analyses.

#### Selection reaction monitoring (SRM) quantitative proteomics

In addition, a small set of targeted proteins (N=119) was measured from post-mortem DLPFC brain tissue of 381 cognitively normal individuals, 286 MCI, 519 AD and 22 with other cause of dementia (68% women) using selection reaction monitoring (SRM) quantitative proteomics as described previously (Yu et al., 2018).

#### RNA-seq

RNA sequencing was performed from DLPFC (N=1102) using a method that has been previously described (Raj et al., 2018). Briefly, samples were extracted using Qiagen’s miRNeasy mini kit (cat. no. 217004) and the RNase free DNase Set (cat. no. 79254), and quantified by Nanodrop and quality was evaluated by Agilent Bioanalyzer. Sequencing was performed on the Illumina HiSeq with 101-bp paired-end reads, the mean coverage for the samples to pass quality control was 95 million reads (median 90 million reads). Data included in the analysis met quality control criteria. For each gene, a normalized expression level was computed by subtracting the mean expression for that gene across all samples and dividing by the SD. The expression levels were adjusted for batch, library size, percentage of coding bases, percentage of aligned reads, percentage of ribosomal bases, percentage of UTR base, median 5 prime to 3 prime base, median CV coverage, post-mortem interval (PMI), and study (ROS or MAP). The data is available through the synapse.org AMP-AD Knowledge Portal (www.synapse.org; SynapseID: syn3388564).

#### Deconvolution of cell type proportion

Proportion of neurons were estimated from DLPFC RNA-seq data by applying the Digital Sorting Algorithm (DSA) (Zhong et al., 2013) to genes previously used to deconvolute cortical RNA-seq data (Wang et al., 2020). RNA-seq data was TMM normalized and technical variables were regressed out. A mean transcription level ≥2 cpm was used to include marker genes of neurons (n=90). The median transcription level of all marker genes per cell type was calculated as described previously by (Klein et al., 2021); Wang et al. (2020).

#### Assessment of DLPFC WGS mtDNAcn

WGS libraries were prepared using the KAPA Hyper Library Preparation Kit in accordance with the manufacturer’s instructions. Briefly, Covaris LE220 sonicator (adaptive focused acoustics) was used to shear 650ng of DNA. Bead-based size selection was performed and selected DNA fragments were then end-repaired, adenylated, and ligated to Illumina sequencing adapters. Fluorescent-based assays including qPCR with the Universal KAPA Library Quantification Kit and Fragment Analyzer (Advanced Analytics) or BioAnalyzer (Agilent 2100) were used to evaluate final libraries. Libraries were sequenced on an Illumina HiSeq X sequencer (v2.5 chemistry) using 2 × 150bp cycles.

R/Bioconductor (packages GenomicAlignments and GenomicRanges) was used to calculate the median sequence coverages of the autosomal chromosomes and of the mitochondrial genome. The intra-contig ambiguity mask from the BSgenome package was used to exclude ambiguous regions. The mtDNAcn was defined as (covmt/covnuc) × 2. the mtDNAcn was z-standardized and then logarithmized as described previously (Klein et al., 2021).

#### Banner AND BLSA LC-MS/MS proteomics

In Banner and BLSA, protein identification and quantification were performed using Liquid Chromatography Coupled to Tandem Mass Spectrometry (LC-MS/MS) as described in (Seyfried et al., 2017; Swarup et al., 2020). Briefly, tissue homogenization and protein digestion were performed. Resulting peptides were desalted with a Sep-Pak C18 column (Waters) and dried under vacuum. Peptide mixtures were separated on a self-packed C18 (1.9 um Dr. Maisch, Germany) fused silica column (25 cm x 75 μM internal diameter; New Objective, Woburn, MA) by a NanoAcquity UHPLC (Waters, Milford, FA) and monitored on a Q-Exactive Plus mass spectrometer (ThermoFisher Scientific, San Jose, CA). MaxQuant (v1.5.2.) with Thermo Foundation 2.0 for RAW file reading capability was used to generate label-free quantification. The quantitation method only considered razor plus unique peptides for protein level quantitation. The protein abundances were log2 transformed and regressed for age at death and postmortem interval.

## QUANTIFICATION AND STATISTICAL ANALYSIS

To assess the extent to which mitochondrial proteins are correlated with each other within RC complexes or between RC we used Spearman correlation matrices. Spearman’s r matrixes were first computed and averaged to compare between groups of protein subunits. The same method was used to assess the cross-talk between nDNA and mtDNA proteins. As well as the overall correlation between RC proteins abundance and mito content, mtDNAcn, mtDNA density and neuron proportion. Before comparing levels of co-regulation between groups of subunits, we tested whether the variables were normally distributed using the Kolmogorov-Smirnov test. For the comparison of two groups, we used either the unpaired t-test or the non-parametric Mann-Whitney test when the variables were not normally distributed. For the comparison of more than two groups, we tested equality of variance using the Brown–Forsyathe test, we used one-way ANOVA with Tukey’s multiple comparison test or the non-parametric equivalent Kruskal-Wallis test with Dunn’s multiple comparisons test when the variables were not normally distributed. Multivariate linear models were used to compare group difference in RC abundance by sex (adjusted for AD status and age at death) and AD status (adjusted by sex and age at death (and years of education in ROSMAP and BLSA)). Effect sizes between groups were computed as Hedges’ g (g) to quantify the magnitude of group differences. Statistical analyses were performed with Prism 8 (GraphPad, CA) and RStudio version 1.3.1056. Statistical significance was set at p<0.05.

### Definition of mitochondrial mass and RC content

Each datapoint was transformed into a z-score (x-mean/sd) before being transformed into a t-score with an average of 100 and a standard deviation of 10 ((x*10)+100).

#### Index of mitochondrial content (mass)

To build a multivariate index of mitochondrial content, we averaged the t-score protein levels of the citrate synthase (CS), the translocator of the outer mitochondrial membrane (TOM20) and voltage dependent anion channel (VDAC1). CS is a matrix-located Krebs cycle enzyme whose enzymatic activity has been validated has a robust marker of mitochondrial mass in human skeletal muscle (Larsen et al., 2012) and has been used as marker of mitochondrial content in the brain (e.g. (Scaini et al., 2010)). TOM20 and VDAC1 are integral proteins of the outer and inner mitochondrial membranes, respectively, and are widely used as a marker of mitochondrial mass (Gouspillou et al., 2014; Narendra et al., 2008; Rehman et al., 2012).

#### Indices of mitochondrial respiratory chain abundance

For each RC complex, t-score protein levels were averaged to build a score representing the average abundance based on all subunits of RC complex I, II, III, IV and V. The proteins included for each complex, and their annotation, are provided in Table S5.

#### Index of mitochondrial DNA density

The ratio of mtDNA copies per cell (mtDNAcn derived from WGS) relative to mitochondrial content per cell (mito content index) was computed to build a score representing mtDNA density per mitochondrion.

## SUPPLEMENTAL ITEM TITLES

### Supplementary figures

Fig. S1. Study design

Fig. S2. Analytical procedure to compute co-regulation.

Fig. S3. Mitochondrial respiratory chain (RC) complexes proteins co-regulation in ROSMAP (TMT).

Fig. S4. Mitochondrial respiratory chain (RC) proteins co-regulation in Banner. Fig. S5. Mitochondrial respiratory chain (RC) proteins co-regulation in BLSA.

Fig. S6. Mitochondrial respiratory chain (RC) proteins co-regulation in ROSMAP. Data from Selected reaction monitoring proteomics (SRM).

Fig. S7. Mitochondrial respiratory chain (RC) transcript co-regulation of the dorsolateral prefrontal cortex (DLPFC).

Fig. S8. Mitochondrial respiratory chain (RC) transcript co-regulation of the posterior cingulate cortex (PCC).

Fig. S7. Mitochondrial respiratory chain (RC) transcript co-regulation of the dorsolateral prefrontal cortex (DLPFC).

Fig. S8. Mitochondrial respiratory chain (RC) transcript co-regulation of the posterior cingulate cortex (PCC).

Fig. S9. Mitochondrial respiratory chain (RC) transcript co-regulation of the anterior caudate (AC).

Fig. S10. Mitochondrial respiratory chain (RC) supercomplexes transcript co-regulation (RNA-seq).

Fig. S11. Mitochondrial respiratory chain (RC) supercomplexes proteins and transcripts co-regulation.

Fig. S12. Nuclear-mtDNA crosstalk by mtDNA subunits.

Fig. S13. Mitochondrial transcription factors (TFs) gene expression and respiratory chain (RC) protein abundance.

Fig. S14. Association between nDNA-encoded mitochondrial respiratory chain (RC) transcript and transcription factors (TFs) levels (RNA-seq).

Fig. S15. Correlations between mitochondrial respiratory chain (RC) proteins (TMT) and transcript (RNAseq) levels in human dorsolateral prefrontal cortex (DLPFC).

Fig. S16. Mitochondrial respiratory chain (RC) proteins co-regulation according to sex and cognitive status.

Fig. S17. Difference in mitochondrial respiratory chain (RC) complex I proteins abundance according to sex and cognitive status.

### Supplementary tables

Supplementary table 1: Participant descriptive statistics

Supplementary table 2: Correlation matrices

Supplementary table 3: Correlation matrices stratified by sex and AD status

Supplementary table 4: Difference in RC protein abundance by sex and AD status

Supplementary table 5: Annotated protein list

## Notes

### Competing Interest Statement

The authors have declared no competing interest.

## References

A Bennett, D., A Schneider, J., Arvanitakis, Z., and S Wilson, R. (2012a). Overview and findings from the religious orders study. Current Alzheimer Research 9, 628–645.

A Bennett, D., A Schneider, J., S Buchman, A., L Barnes, L., A Boyle, P., and S Wilson, R. (2012b). Overview and findings from the rush Memory and Aging Project. Current Alzheimer Research 9, 646–663.

Acín-Pérez, R., Fernández-Silva, P., Peleato, M.L., Pérez-Martos, A., and Enriquez, J.A. (2008). Respiratory active mitochondrial supercomplexes. Molecular cell 32, 529–539.

Beach, T.G., Adler, C.H., Sue, L.I., Serrano, G., Shill, H.A., Walker, D.G., Lue, L., Roher, A.E., Dugger, B.N., and Maarouf, C. (2015). Arizona study of aging and neurodegenerative disorders and brain and body donation program. Neuropathology: official journal of the Japanese Society of Neuropathology 35, 354.

Bennett, D., Schneider, J., Arvanitakis, Z., Kelly, J., Aggarwal, N., Shah, R., and Wilson, R. (2006). Neuropathology of older persons without cognitive impairment from two community-based studies. Neurology 66, 1837–1844.

Bennett, D., Wilson, R., Schneider, J., Evans, D., Aggarwal, N., Arnold, S., Cochran, E., Berry-Kravis, E., and Bienias, J. (2003). Apolipoprotein E ε4 allele, AD pathology, and the clinical expression of Alzheimer’s disease. Neurology 60, 246–252.

Bennett, D.A., Buchman, A.S., Boyle, P.A., Barnes, L.L., Wilson, R.S., and Schneider, J.A. (2018). Religious orders study and rush memory and aging project. Journal of Alzheimer’s Disease 64, S161–S189.

Chung-ha, O.D., Kim, K.-Y., Bushong, E.A., Mills, E.A., Boassa, D., Shih, T., Kinebuchi, M., Phan, S., Zhou, Y., and Bihlmeyer, N.A. (2014). Transcellular degradation of axonal mitochondria. Proceedings of the National Academy of Sciences 111, 9633–9638.

de Sousa Abreu, R., Penalva, L.O., Marcotte, E.M., and Vogel, C. (2009). Global signatures of protein and mRNA expression levels. Molecular BioSystems 5, 1512–1526.

Ferrucci, L. (2008). The Baltimore Longitudinal Study of Aging (BLSA): a 50-year-long journey and plans for the future (Oxford University Press).

Filograna, R., Mennuni, M., Alsina, D., and Larsson, N.G. (2020). Mitochondrial DNA copy number in human disease: the more the better? FEBS letters.

Formosa, L.E., Dibley, M.G., Stroud, D.A., and Ryan, M.T. (2018). Building a complex complex: assembly of mitochondrial respiratory chain complex I. Paper presented at: Seminars in Cell & Developmental Biology (Elsevier).

Garcia, C.J., Khajeh, J., Coulanges, E., Emily, I., and Owusu-Ansah, E. (2017). Regulation of mitochondrial complex I biogenesis in Drosophila flight muscles. Cell reports 20, 264–278.

Giordano, C., Iommarini, L., Giordano, L., Maresca, A., Pisano, A., Valentino, M.L., Caporali, L., Liguori, R., Deceglie, S., and Roberti, M. (2014). Efficient mitochondrial biogenesis drives incomplete penetrance in Leber’s hereditary optic neuropathy. Brain 137, 335–353.

Gómez-Isla, T., Hollister, R., West, H., Mui, S., Growdon, J.H., Petersen, R.C., Parisi, J.E., and Hyman, B.T. (1997). Neuronal loss correlates with but exceeds neurofibrillary tangles in Alzheimer’s disease. Annals of Neurology: Official Journal of the American Neurological Association and the Child Neurology Society 41, 17–24.

Gouspillou, G., Sgarioto, N., Norris, B., Barbat-Artigas, S., Aubertin-Leheudre, M., Morais, J.A., Burelle, Y., Taivassalo, T., and Hepple, R.T. (2014). The relationship between muscle fiber type-specific PGC-1α content and mitochondrial content varies between rodent models and humans. PloS one 9, e103044.

Grünewald, A., Rygiel, K.A., Hepplewhite, P.D., Morris, C.M., Picard, M., and Turnbull, D.M. (2016). Mitochondrial DNA Depletion in Respiratory Chain–Deficient P arkinson Disease Neurons. Annals of neurology 79, 366–378.

Holper, L., Ben-Shachar, D., and Mann, J. (2019). Multivariate meta-analyses of mitochondrial complex I and IV in major depressive disorder, bipolar disorder, schizophrenia, Alzheimer disease, and Parkinson disease. Neuropsychopharmacology 44, 837–849.

Hou, T., Zhang, R., Jian, C., Ding, W., Wang, Y., Ling, S., Ma, Q., Hu, X., Cheng, H., and Wang, X. (2019). NDUFAB1 confers cardio-protection by enhancing mitochondrial bioenergetics through coordination of respiratory complex and supercomplex assembly. Cell research 29, 754–766.

Houtkooper, R.H., Mouchiroud, L., Ryu, D., Moullan, N., Katsyuba, E., Knott, G., Williams, R.W., and Auwerx, J. (2013). Mitonuclear protein imbalance as a conserved longevity mechanism. Nature 497, 451–457.

Hyman, B.T., and Trojanowski, J.Q. (1997). Editorial on consensus recommendations for the postmortem diagnosis of Alzheimer disease from the National Institute on Aging and the Reagan Institute Working Group on diagnostic criteria for the neuropathological assessment of Alzheimer disease. Journal of neuropathology and experimental neurology 56, 1095.

Jiang, L., Wang, M., Lin, S., Jian, R., Li, X., Chan, J., Dong, G., Fang, H., Robinson, A.E., and Aguet, F. (2020). A quantitative proteome map of the human body. Cell 183, 269-283. e219.

Johnson, E.C., Dammer, E.B., Duong, D.M., Yin, L., Thambisetty, M., Troncoso, J.C., Lah, J.J., Levey, A.I., and Seyfried, N.T. (2018). Deep proteomic network analysis of Alzheimer’s disease brain reveals alterations in RNA binding proteins and RNA splicing associated with disease. Molecular neurodegeneration 13, 1–22.

Kim, K.H., Son, J.M., Benayoun, B.A., and Lee, C. (2018). The mitochondrial-encoded peptide MOTS-c translocates to the nucleus to regulate nuclear gene expression in response to metabolic stress. Cell metabolism 28, 516-524. e517.

Klein, H.-U., Schäfer, M., Bennett, D.A., Schwender, H., and De Jager, P.L. (2020). Bayesian integrative analysis of epigenomic and transcriptomic data identifies Alzheimer’s disease candidate genes and networks. PLoS computational biology 16, e1007771.

Klein, H.-U., Trumpff, C., Yang, H.-S., Lee, A.J., Picard, M., Bennett, D.A., and De Jager, P.L. (2021). Characterization of mitochondrial DNA quantity and quality in the human aged and Alzheimer’s disease brain. medRxiv, 2021.2005.2020.21257456.

Kong, D., Zhao, L., Du, Y., He, P., Zou, Y., Yang, L., Sun, L., Wang, H., Xu, D., and Meng, X. (2014). Overexpression of GRIM-19, a mitochondrial respiratory chain complex I protein, suppresses hepatocellular carcinoma growth. International journal of clinical and experimental pathology 7, 7497.

Kummer, E., and Ban, N. (2021). Mechanisms and regulation of protein synthesis in mitochondria. Nature Reviews Molecular Cell Biology, 1–19.

Kustatscher, G., Grabowski, P., Schrader, T.A., Passmore, J.B., Schrader, M., and Rappsilber, J. (2019). Co-regulation map of the human proteome enables identification of protein functions. Nature biotechnology 37, 1361–1371.

Lapuente-Brun, E., Moreno-Loshuertos, R., Acín-Pérez, R., Latorre-Pellicer, A., Colás, C., Balsa, E., Perales-Clemente, E., Quirós, P.M., Calvo, E., and Rodríguez-Hernández, M. (2013). Supercomplex assembly determines electron flux in the mitochondrial electron transport chain. Science 340, 1567–1570.

Larsen, S., Nielsen, J., Hansen, C.N., Nielsen, L.B., Wibrand, F., Stride, N., Schroder, H.D., Boushel, R., Helge, J.W., and Dela, F. (2012). Biomarkers of mitochondrial content in skeletal muscle of healthy young human subjects. The Journal of physiology 590, 3349–3360.

Letts, J.A., and Sazanov, L.A. (2017). Clarifying the supercomplex: the higher-order organization of the mitochondrial electron transport chain. Nature structural & molecular biology 24, 800–808.

Maier, T., Güell, M., and Serrano, L. (2009). Correlation of mRNA and protein in complex biological samples. FEBS letters 583, 3966–3973.

McLaughlin, K.L., Hagen, J.T., Coalson, H.S., Nelson, M.A., Kew, K.A., Wooten, A.R., and Fisher-Wellman, K.H. (2020). Novel approach to quantify mitochondrial content and intrinsic bioenergetic efficiency across organs. Scientific reports 10, 1–15.

Mertins, P., Tang, L.C., Krug, K., Clark, D.J., Gritsenko, M.A., Chen, L., Clauser, K.R., Clauss, T.R., Shah, P., and Gillette, M.A. (2018). Reproducible workflow for multiplexed deep-scale proteome and phosphoproteome analysis of tumor tissues by liquid chromatography–mass spectrometry. Nature protocols 13, 1632–1661.

Milenkovic, D., Blaza, J.N., Larsson, N.-G., and Hirst, J. (2017). The enigma of the respiratory chain supercomplex. Cell metabolism 25, 765–776.

Mittelstrass, K., Ried, J.S., Yu, Z., Krumsiek, J., Gieger, C., Prehn, C., Roemisch-Margl, W., Polonikov, A., Peters, A., and Theis, F.J. (2011). Discovery of sexual dimorphisms in metabolic and genetic biomarkers. PLoS genetics 7, e1002215.

Mostafavi, S., Gaiteri, C., Sullivan, S.E., White, C.C., Tasaki, S., Xu, J., Taga, M., Klein, H.-U., Patrick, E., and Komashko, V. (2018). A molecular network of the aging human brain provides insights into the pathology and cognitive decline of Alzheimer’s disease. Nature neuroscience 21, 811–819.

Mukherjee, S., and Ghosh, A. (2020). Molecular mechanism of mitochondrial respiratory chain assembly and its relation to mitochondrial diseases. Mitochondrion.

Narendra, D., Tanaka, A., Suen, D.-F., and Youle, R.J. (2008). Parkin is recruited selectively to impaired mitochondria and promotes their autophagy. The Journal of cell biology 183, 795–803.

Nicholls, D.G. (2013). Bioenergetics (Academic Press).

Picard, M., and McEwen, B.S. (2014). Mitochondria impact brain function and cognition. Proceedings of the National Academy of Sciences 111, 7–8.

Ping, L., Duong, D.M., Yin, L., Gearing, M., Lah, J.J., Levey, A.I., and Seyfried, N.T. (2018). Global quantitative analysis of the human brain proteome in Alzheimer’s and Parkinson’s Disease. Scientific data 5, 1–12.

Quirós, P.M., Mottis, A., and Auwerx, J. (2016). Mitonuclear communication in homeostasis and stress. Nature reviews Molecular cell biology 17, 213.

Raj, T., Li, Y.I., Wong, G., Humphrey, J., Wang, M., Ramdhani, S., Wang, Y.-C., Ng, B., Gupta, I., and Haroutunian, V. (2018). Integrative transcriptome analyses of the aging brain implicate altered splicing in Alzheimer’s disease susceptibility. Nature genetics 50, 1584–1592.

Rangaraju, V., Lewis, T.L., Hirabayashi, Y., Bergami, M., Motori, E., Cartoni, R., Kwon, S.-K., and Courchet, J. (2019). Pleiotropic mitochondria: the influence of mitochondria on neuronal development and disease. Journal of Neuroscience 39, 8200–8208.

Rath, S., Sharma, R., Gupta, R., Ast, T., Chan, C., Durham, T.J., Goodman, R.P., Grabarek, Z., Haas, M.E., and Hung, W.H. (2020). MitoCarta3. 0: an updated mitochondrial proteome now with sub-organelle localization and pathway annotations. Nucleic Acids Research.

Rehman, J., Zhang, H.J., Toth, P.T., Zhang, Y., Marsboom, G., Hong, Z., Salgia, R., Husain, A.N., Wietholt, C., and Archer, S.L. (2012). Inhibition of mitochondrial fission prevents cell cycle progression in lung cancer. The FASEB Journal 26, 2175–2186.

Robins, C., Wingo, A.P., Fan, W., Doung, D.M., Meigs, J., Gerasimov, E.S., Dammer, E.B., Cutler, D.J., De Jager, P., and Bennett, D.A. (2019). Genetic control of the human brain proteome. bioRxiv, 816652.

Scaini, G., Santos, P.M., Benedet, J., Rochi, N., Gomes, L.M., Borges, L.S., Rezin, G.T., Pezente, D.P., Quevedo, J., and Streck, E.L. (2010). Evaluation of Krebs cycle enzymes in the brain of rats after chronic administration of antidepressants. Brain research bulletin 82, 224–227.

Seyfried, N.T., Dammer, E.B., Swarup, V., Nandakumar, D., Duong, D.M., Yin, L., Deng, Q., Nguyen, T., Hales, C.M., and Wingo, T. (2017). A multi-network approach identifies protein-specific co-expression in asymptomatic and symptomatic Alzheimer’s disease. Cell systems 4, 60-72. e64.

Swarup, V., Chang, T.S., Duong, D.M., Dammer, E.B., Dai, J., Lah, J.J., Johnson, E.C., Seyfried, N.T., Levey, A.I., and Geschwind, D.H. (2020). Identification of conserved proteomic networks in neurodegenerative dementia. Cell Reports 31, 107807.

Swerdlow, R.H., Burns, J.M., and Khan, S.M. (2014). The Alzheimer’s disease mitochondrial cascade hypothesis: progress and perspectives. Biochimica et Biophysica Acta (BBA)-Molecular Basis of Disease 1842, 1219–1231.

Szczepanowska, K., Senft, K., Heidler, J., Herholz, M., Kukat, A., Höhne, M.N., Hofsetz, E., Becker, C., Kaspar, S., and Giese, H. (2020). A salvage pathway maintains highly functional respiratory complex I. Nature communications 11, 1–18.

Szczepanowska, K., and Trifunovic, A. (2021). Tune instead of destroy: How proteolysis keeps OXPHOS in shape. Biochimica et Biophysica Acta (BBA)-Bioenergetics, 148365.

Vartak, R., Porras, C.A.-M., and Bai, Y. (2013). Respiratory supercomplexes: structure, function and assembly. Protein & cell 4, 582–590.

Ventura-Clapier, R., Moulin, M., Piquereau, J., Lemaire, C., Mericskay, M., Veksler, V., and Garnier, A. (2017). Mitochondria: a central target for sex differences in pathologies. Clinical Science 131, 803–822.

Vincent, A.E., Rosa, H.S., Pabis, K., Lawless, C., Chen, C., Grünewald, A., Rygiel, K.A., Rocha, M.C., Reeve, A.K., and Falkous, G. (2018). Subcellular origin of mitochondrial DNA deletions in human skeletal muscle. Annals of neurology 84, 289–301.

Wang, X., Allen, M., Li, S., Quicksall, Z.S., Patel, T.A., Carnwath, T.P., Reddy, J.S., Carrasquillo, M.M., Lincoln, S.J., Nguyen, T.T., et al. (2020). Deciphering cellular transcriptional alterations in Alzheimer’s disease brains. Mol Neurodegener 15, 38.

Wingo, A.P., Dammer, E.B., Breen, M.S., Logsdon, B.A., Duong, D.M., Troncosco, J.C., Thambisetty, M., Beach, T.G., Serrano, G.E., and Reiman, E.M. (2019). Large-scale proteomic analysis of human brain identifies proteins associated with cognitive trajectory in advanced age. Nature communications 10, 1–14.

Yu, L., Petyuk, V.A., Gaiteri, C., Mostafavi, S., Young-Pearse, T., Shah, R.C., Buchman, A.S., Schneider, J.A., Piehowski, P.D., and Sontag, R.L. (2018). Targeted brain proteomics uncover multiple pathways to Alzheimer’s dementia. Annals of neurology 84, 78–88.

Yu, L., Tasaki, S., Schneider, J.A., Arfanakis, K., Duong, D.M., Wingo, A.P., Wingo, T.S., Kearns, N., Thatcher, G.R., and Seyfried, N.T. (2020). Cortical proteins associated with cognitive resilience in community-dwelling older persons. JAMA psychiatry 77, 1172–1180.

Yu-Wai-Man, P., Sitarz, K.S., Samuels, D.C., Griffiths, P.G., Reeve, A.K., Bindoff, L.A., Horvath, R., and Chinnery, P.F. (2010). OPA1 mutations cause cytochrome c oxidase deficiency due to loss of wild-type mtDNA molecules. Human molecular genetics 19, 3043–3052.

Zhang, R., Hou, T., Cheng, H., and Wang, X. (2019). NDUFAB1 protects against obesity and insulin resistance by enhancing mitochondrial metabolism. The FASEB Journal 33, 13310–13322.

Zhong, Y., Wan, Y.W., Pang, K., Chow, L.M., and Liu, Z. (2013). Digital sorting of complex tissues for cell type-specific gene expression profiles. BMC Bioinformatics 14, 89.

