## Supplemental figures for "Mitochondrial Respiratory Chain Protein Co-Regulation in the Human Brain"

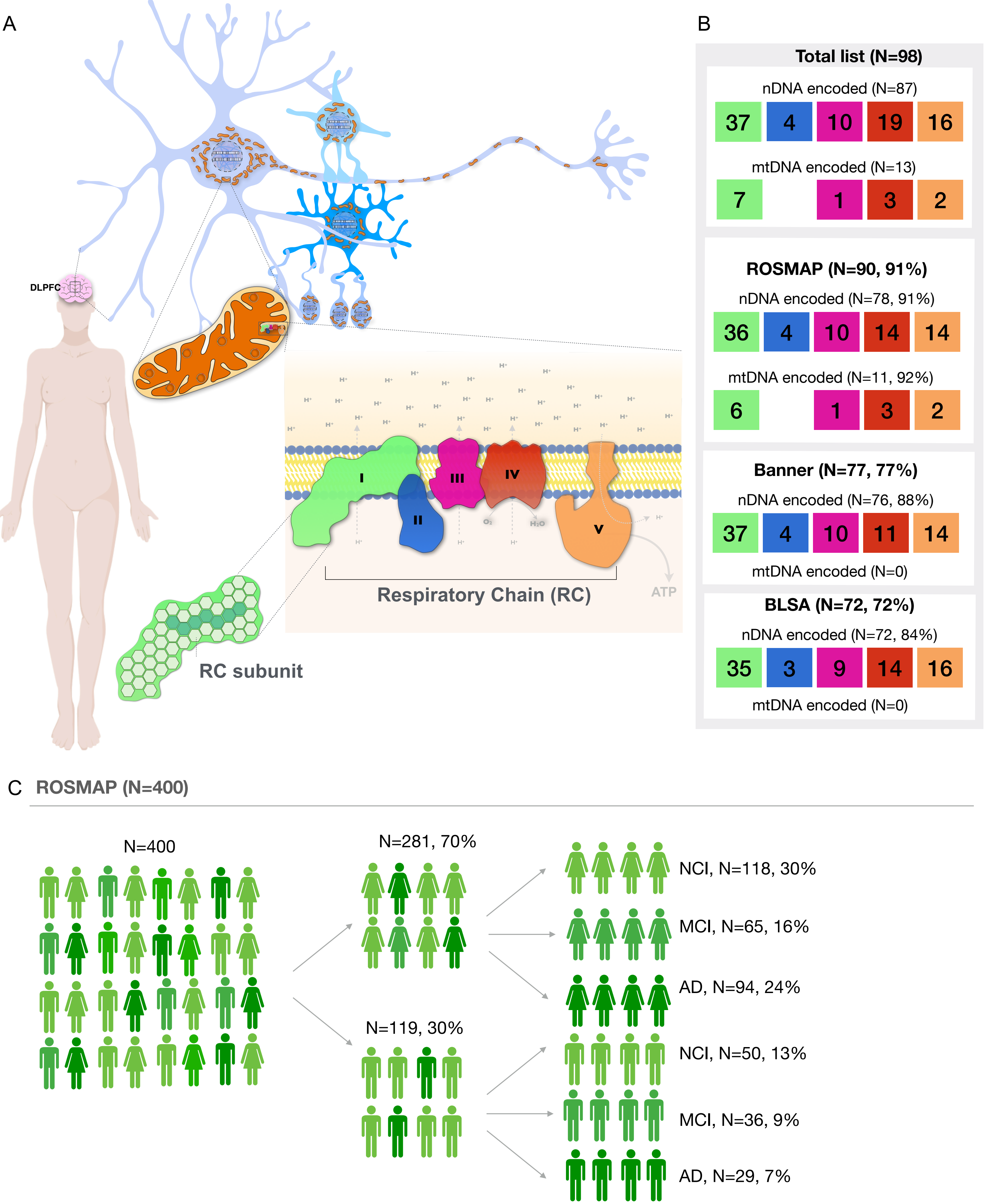


**Fig. S1. Study design** (**A**) co-regulation of mitochondrial respiratory chain (RC) proteins (or subunits) in the human dorsolateral prefrontal cortex (DLPFC) was assessed across three independent cohorts: ROSMAP (N=400), BLSA (N=47) and Banner (N=201). (**B**) Coverage of gene encoding for RC proteins by study cohort. Participants in ROSMAP were not cognitively impaired (NCI) at enrollment, some of them developed mild cognitive impairment (MCI) or Alzheimer disease (AD). (**C**) ROSMAP participants characteristics for which proteomic data is available. Detailed participants characteristics for ROSMAP (N=400), BLSA (N=47) and Banner (N=201) are shown in supplementary table S1.

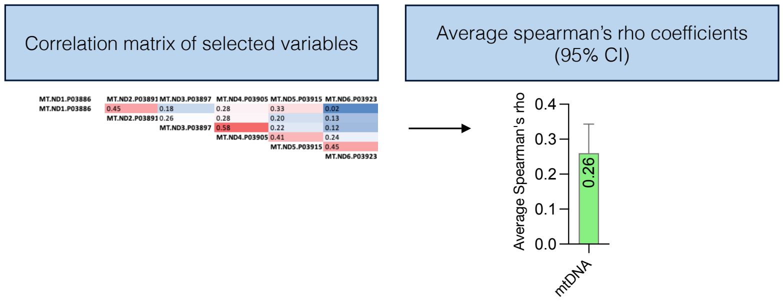


**Fig. S2. Analytical procedure to compute co-regulation.** Example of computation for mtDNA encoded complex I subunits co-regulation.

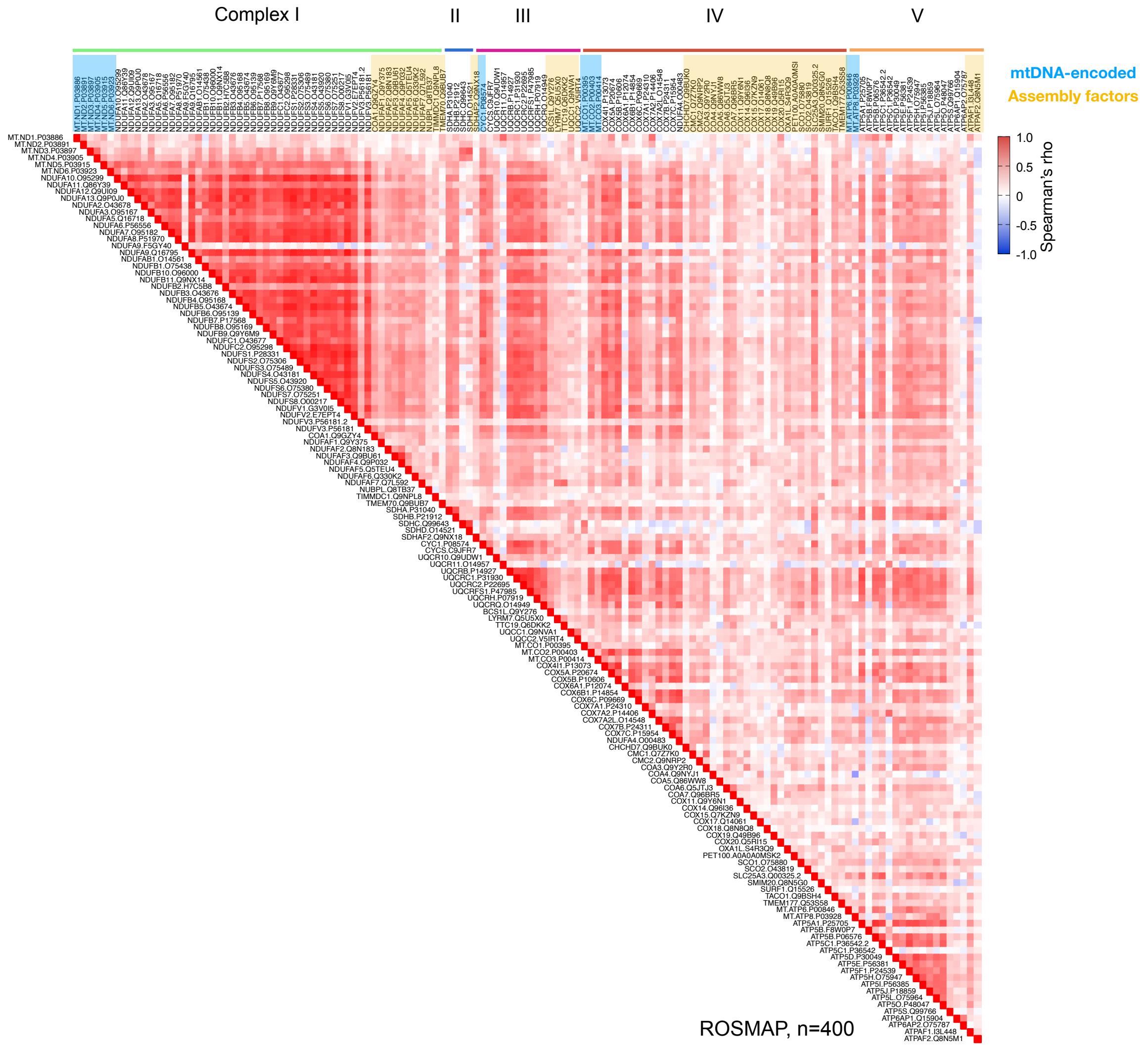


**Fig. S3. Mitochondrial respiratory chain (RC) complexes proteins co-regulation in ROSMAP (TMT).** (**A**) Heatmap of RC subunits and assembly factors associations assessed with Spearman rank correlation. ROSMAP, N=400. Correlation matrix is shown in supplemental table S2A.

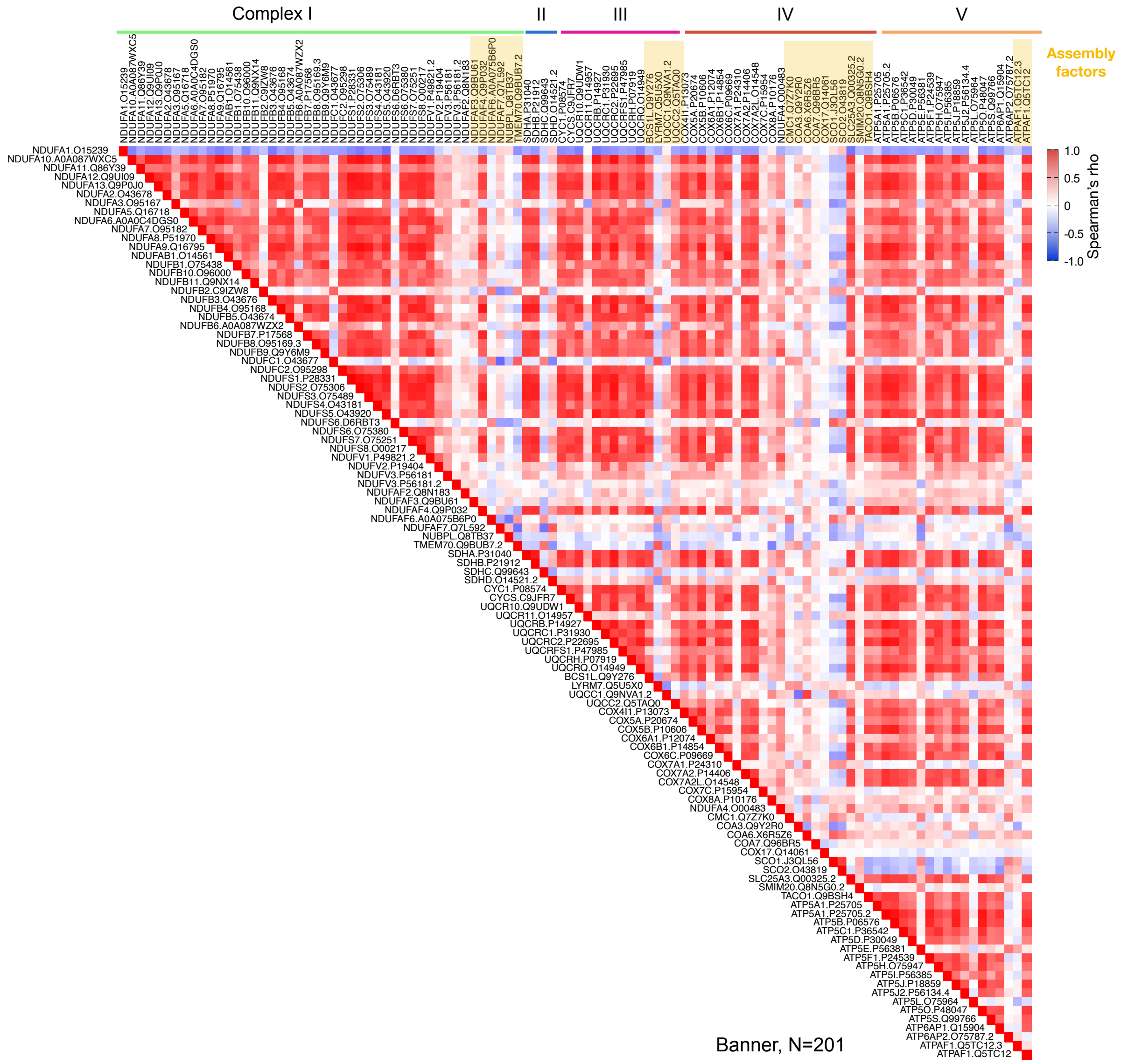


**Fig. S4. Mitochondrial respiratory chain (RC) proteins co-regulation in Banner.** (**A**) Heatmap of RC subunits and assembly factors associations assessed with Spearman rank correlation. Banner, N=201. Correlation matrix is shown in supplemental table S2B.

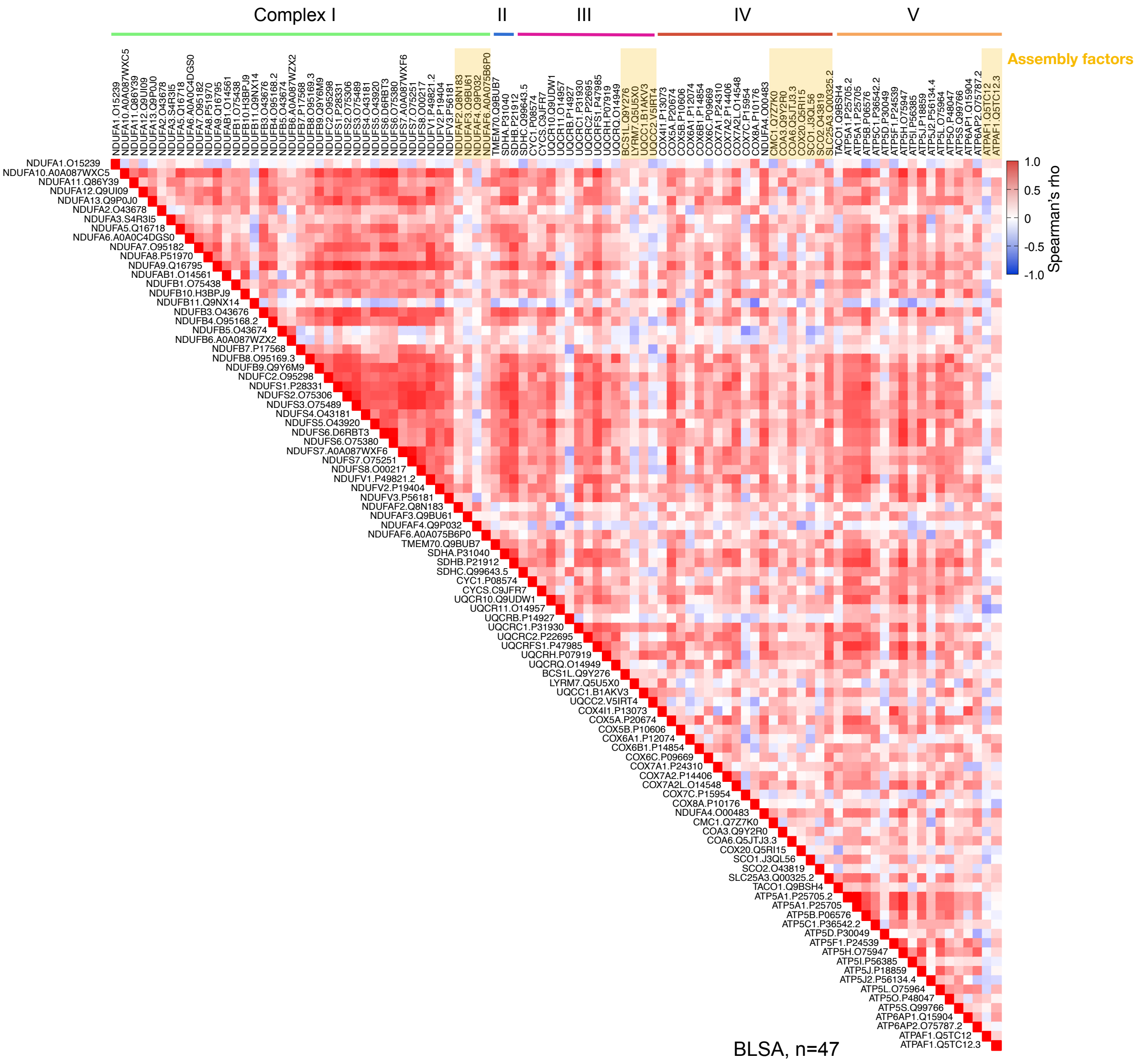


**Fig. S5. Mitochondrial respiratory chain (RC) proteins co-regulation in BLSA.** (**A**) Heatmap of RC subunits and assembly factors associations assessed with Spearman rank correlation. BLSA, N=47. Correlation matrix is shown in supplemental table S2C.

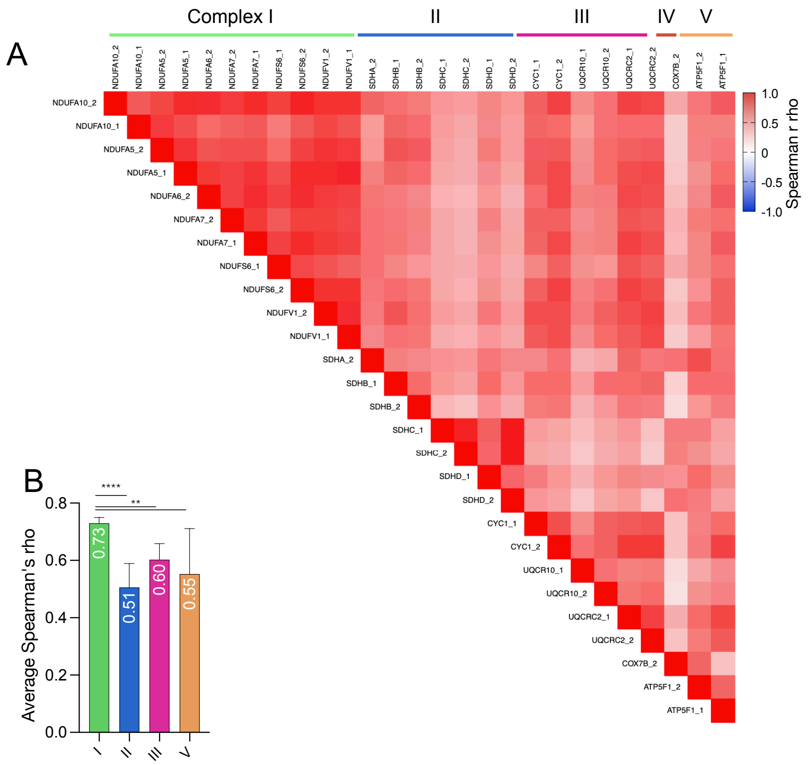


**Fig. S6. Mitochondrial respiratory chain (RC) proteins co-regulation in ROSMAP. Data from Selected reaction monitoring proteomics (SRM).** (**A**) Heatmap of nuclear DNA (nDNA) encoded RC subunits associations assessed with spearman rank correlation. (**B**) Average spearman's rho (95% CI) for nDNA encoded RC subunits of complexes I, II, III, V. ROSMAP, N=1228 (388 men, 823 women). Correlation matrix is shown in supplemental table S2D. P-values from One-way ANOVA and Tukey's multiple comparison test, p<0.05*, p<0.01**, p<0.001***, p<0.0001****.

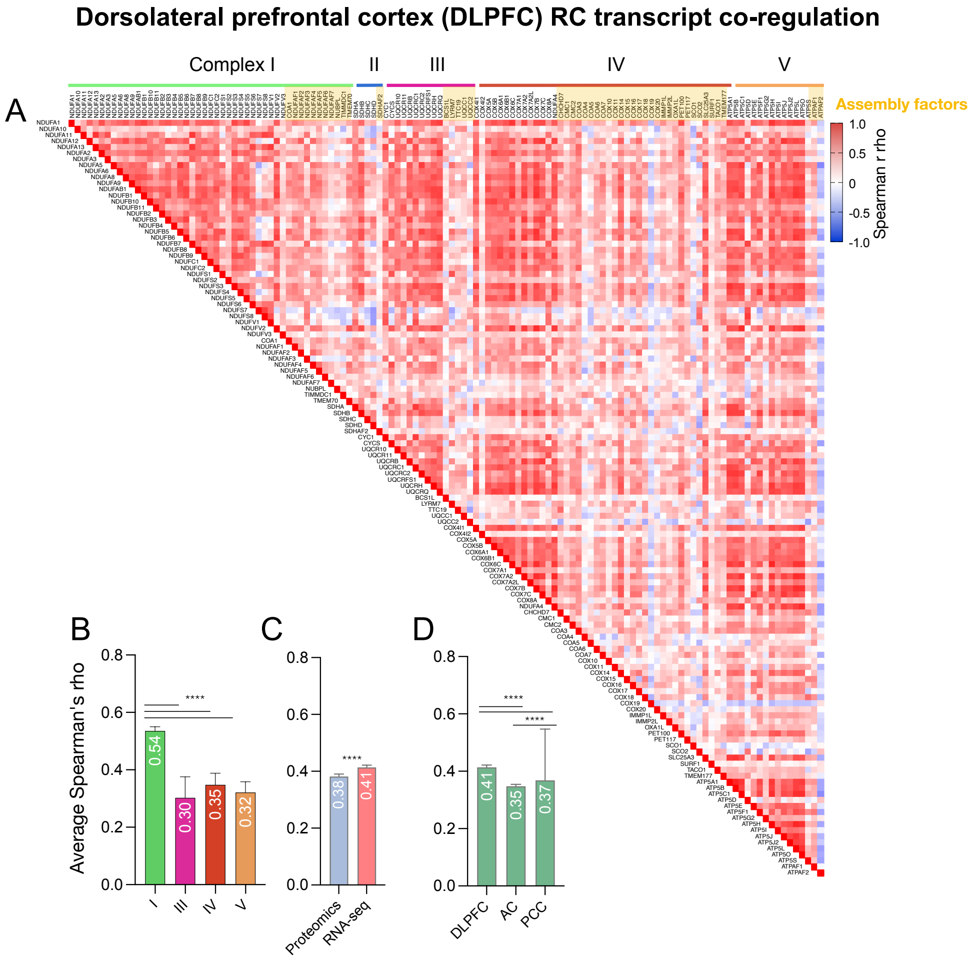


**Fig. S7. Mitochondrial respiratory chain (RC) transcript co-regulation of the dorsolateral prefrontal cortex (DLPFC).** (**A**) Heatmap of RC subunits and assembly factors transcript (RNA-seq) associations assessed with spearman rank correlation. (**B**) Average spearman's rho (95% CI) for nuclear DNA (nDNA) encoded RMC transcripts for complexes I, III, IV, V**.** N = 1092 (**C**) Comparison of average spearman's rho (95% CI) for nDNA encoded RC proteins and transcripts. (**D**) Comparison of average spearman's rho (95% CI) for nDNA encoded RC transcripts in DLPFC, posterior cingulate cortex (PCC) (N=661) and anterior caudate (AC) (N=731). Correlation matrix are shown in supplemental table S2E-G. P-values from Mann-Whitney test or Kruskal-Wallis test with Dunn's multiple comparisons test, p<0.05*, p<0.01**, p<0.001***, p<0.0001****.

### Posterior cingulate cortex (PCC) RC transcript co-regulation

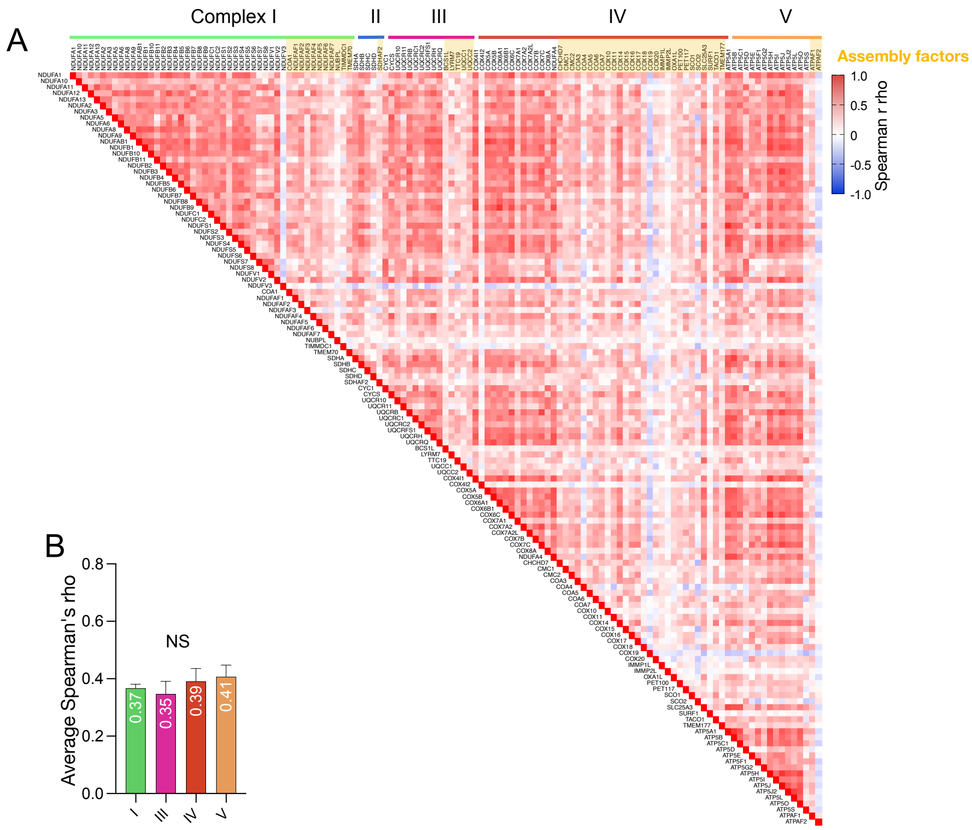


**Fig. S8. Mitochondrial respiratory chain (RC) transcript co-regulation of the posterior cingulate cortex (PCC).** (**A**) Heatmap of RC subunits and assembly factors transcript (RNA-seq) associations assessed with spearman rank correlation. (**B**) Average spearman's rho for nuclear DNA (nDNA) encoded RC transcripts for complexes I, III, IV, V. P-value from Kruskal-Wallis test. N = 661. Correlation matrix is shown in supplemental table S2F.

### Anterior caudate (AC) RC transcript co-regulation

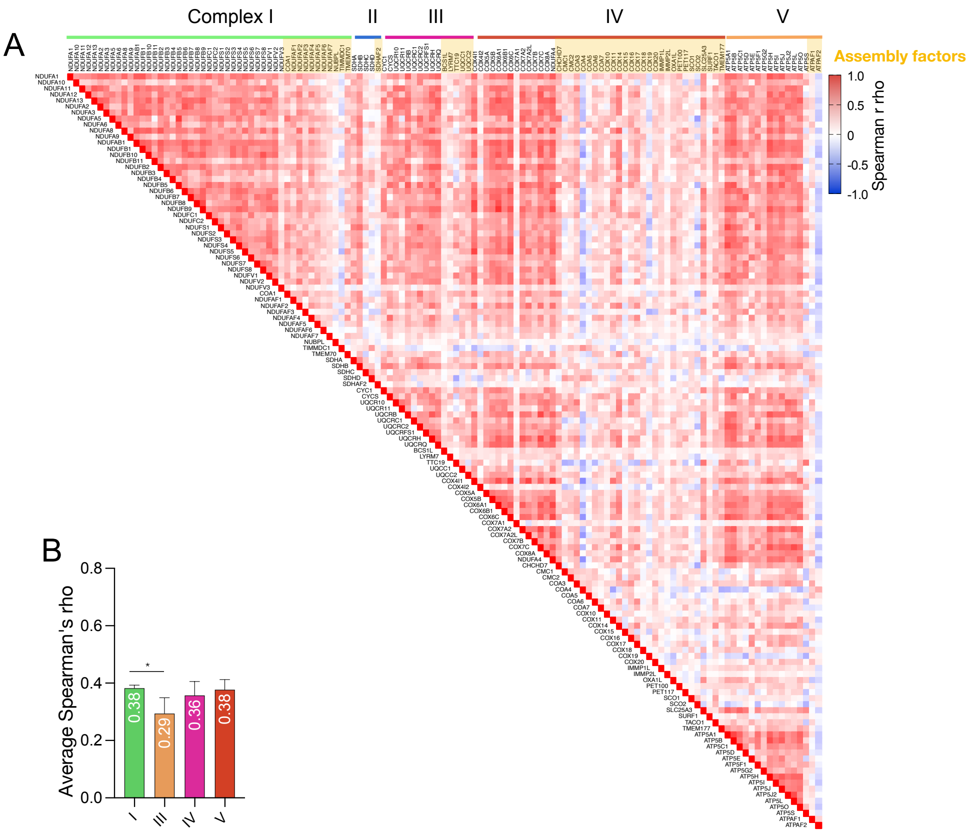


**Fig. S9. Mitochondrial respiratory chain (RC) transcript co-regulation of the anterior caudate (AC).** (**A**) Heatmap of RC subunits and assembly factors transcript (RNA-seq) associations assessed with spearman rank correlation. (**B**) Average spearman's rho (95% CI) for nuclear DNA (nDNA) encoded RC transcripts for complexes I, III, IV, V. N = 731. Correlation matrix is shown in supplemental table S2G. P-values from Kruskal-Wallis test with Dunn's multiple comparisons test, $p<0.05$*, $p<0.01$**, $p<0.001$***, $p<0.0001$****.

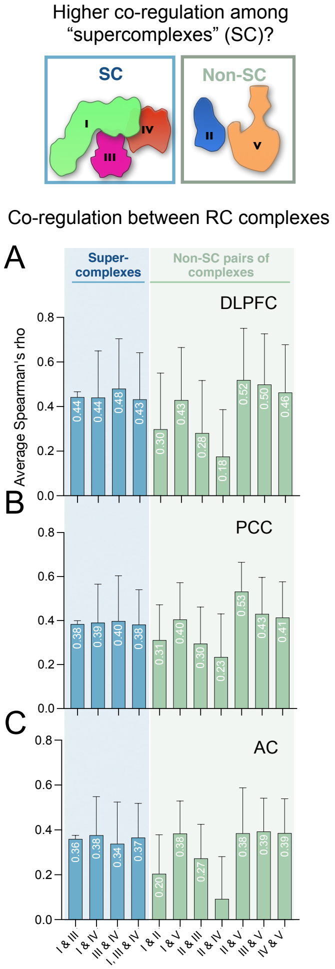


**Fig. S10. Mitochondrial respiratory chain (RC) supercomplexes transcript co-regulation (RNA-seq).** Average spearman's rho (95% CI) for RC complexes part of the hypothesized supercomplexes (SC, I, III, IV) and other possible pairs of RC complexes in the (**A**) dorsolateral prefrontal cortex DLPFC, (**B**) posterior cingulate cortex (PCC), and (**C**) anterior caudate (AC). P-values from Dunn's multiple comparisons test are shown in Fig S11D-F.

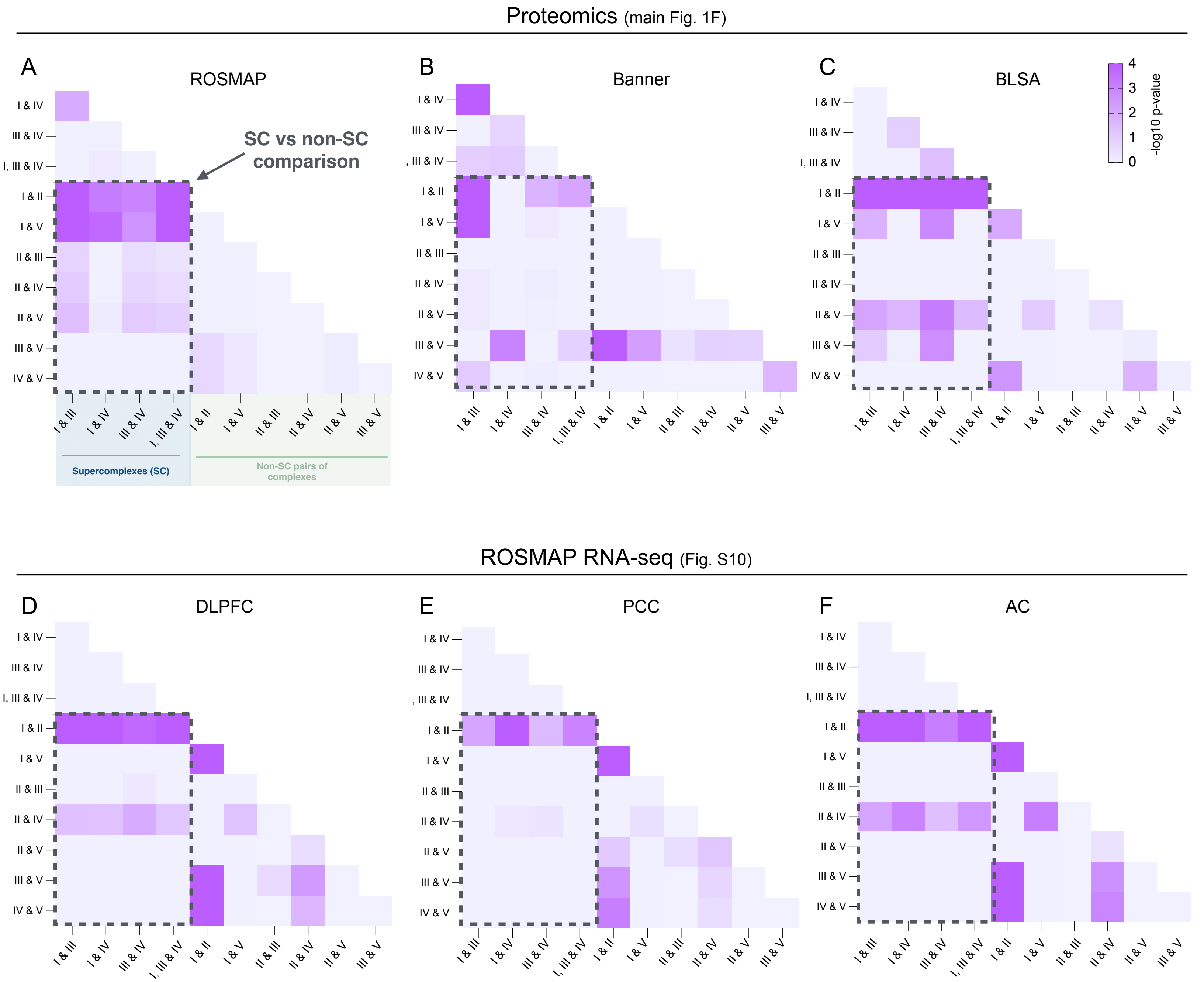


**Fig. S11. Statistics for co-regulation of mitochondrial respiratory chain supercomplexes (SC) proteins and transcripts.** P-values for the difference between average Spearman's rho for RC complexes part of the hypothesized supercomplexes (SC, I, III, IV) and other possible pairs of RC complexes. Results for the dorsolateral prefrontal cortex (DLPFC) proteins in (**A**) ROSMAP (N=400), (**B**) Banner (N=201), (**C**) BLSA (N=47), in reference to results shown in main Fig. 1F. Results for ROSMAP transcripts (RNA-seq) in (**D**) the DLPFC, (**E**) the posterior cingulate cortex (PCC) and (**F**) the anterior caudate (AC). P-values from Dunn's multiple comparisons test.

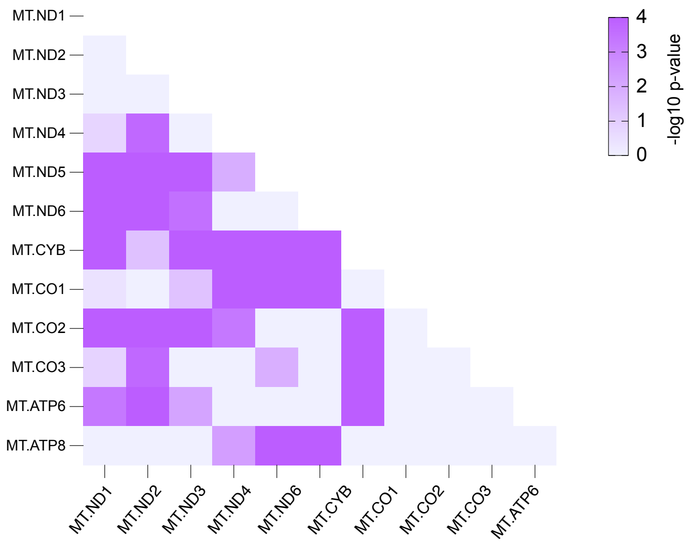


**Fig. S12. Statistics for the co-regulation of mtDNA subunits.** P-values comparing the abundance of mtDNA-encoded subunits from main Fig. 3E. P-values from Dunn's multiple comparisons test.

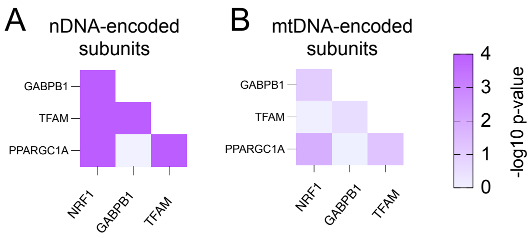


**Fig. S13. Statistics for the mitochondrial transcription factors (TFs) gene expression and respiratory chain (RC) protein abundance.** P-value for the difference in the strength of the association (Spearman's rho) between RC subunits protein abundance and TFs gene expression. Results for (**A**) nuclear DNA (nDNA) and (**B**) mtDNA-encoded subunits (see Fig3H). ROSMAP, N =124-400. P-values from Dunn's multiple comparisons test (A) and Tukey's multiple comparisons test (B).

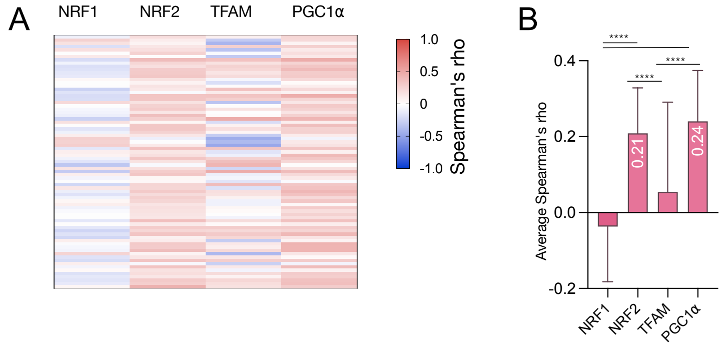


**Fig. S14. Association between nuclear DNA (nDNA) encoded mitochondrial respiratory chain (RC) transcript and transcription factors (TFs) levels (RNA-seq).** (**A**) Heatmap and (**B**) average Spearman's rho (95% CI) of RC subunits transcript abundance and transcription factor gene expression (NRF2 corresponds to GABPB1 gene). Correlation matrix is shown in table S2. P-values from Kruskal-Wallis test with Dunn's multiple comparisons test. ROSMAP, N =1100.

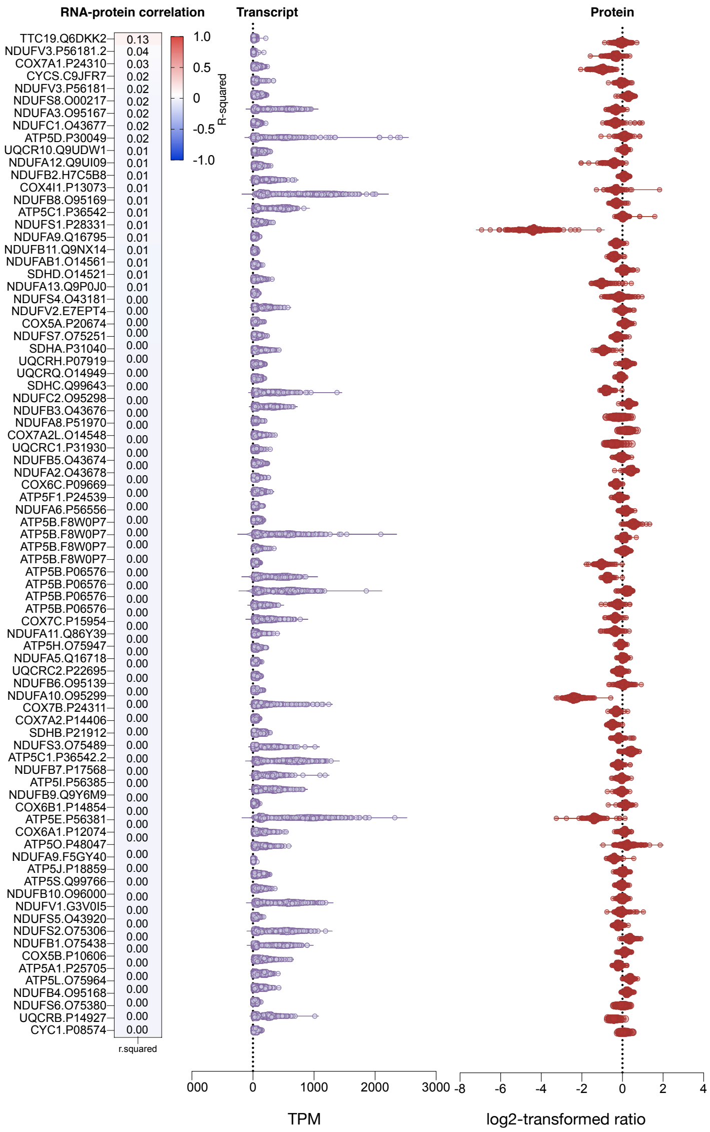

**Fig. S15. Correlations between mitochondrial respiratory chain (RC) proteins (TMT) and transcript (RNAseq) levels in human dorsolateral prefrontal cortex (DLPFC).** (**A**) Heatmap sorted by proportion of shared variance ($r^2$, from linear regression). Raw data (unadjusted) was used for this analysis. ROSMAP, N=124-400.

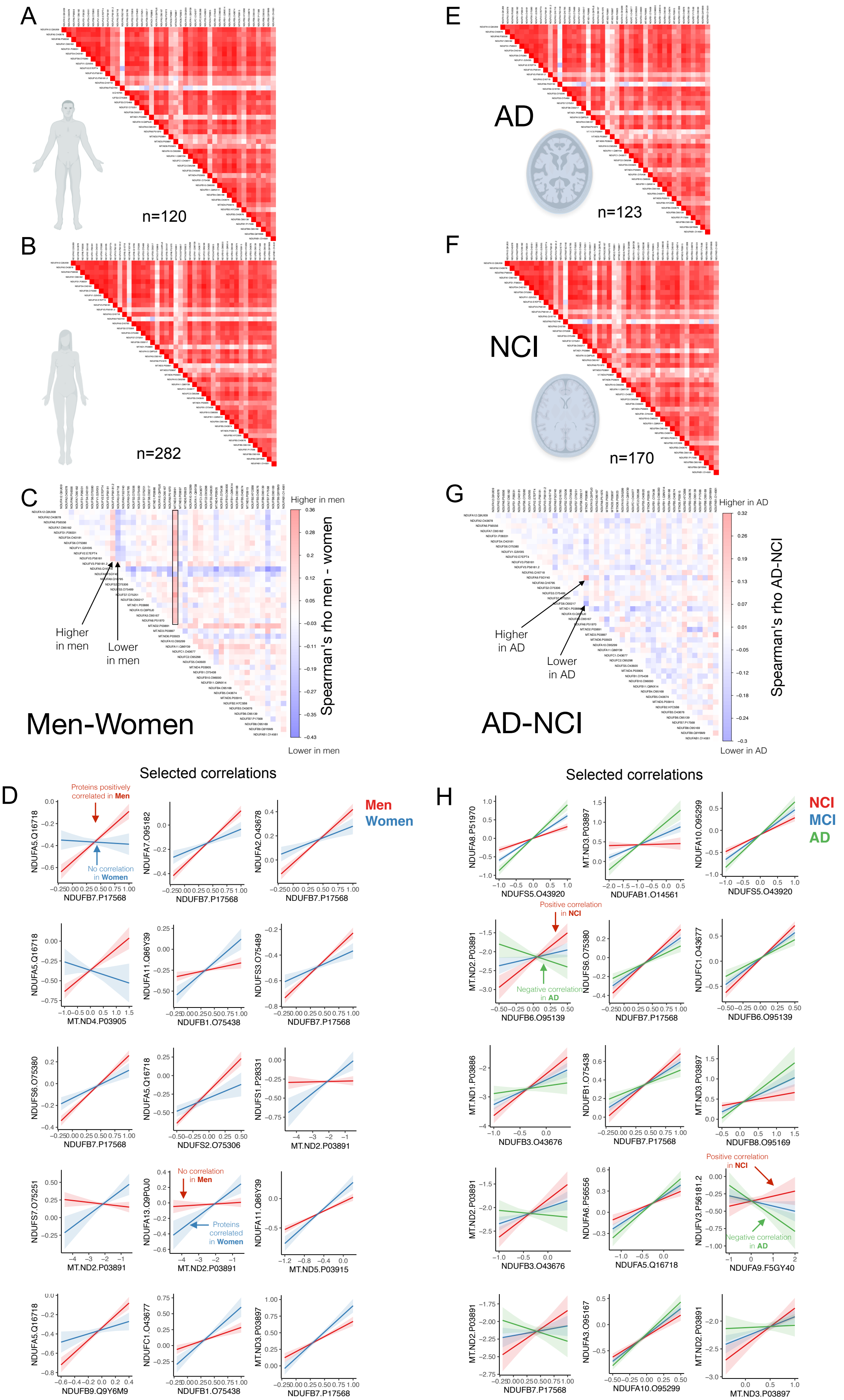


**Fig. S16. Mitochondrial respiratory chain (RC) proteins co-regulation according to sex and clinical diagnosis.** Complex subunits co-regulation for (**A**) men (**B**) women. (**C**) Difference in Spearman's rho coefficients between men and women. The correlations that differ between the sexes appear in darker colors. (**D**) Interaction represented for the top 15 most significant interactions computed on the 100 most different pairs of subunits. Complex subunits co-regulation for (**E**) individuals with Alzheimer's disease (AD) or (**F**) non-cognitively impaired (NCI). (**G**) Difference in Spearman's rho coefficients between AD and NCI. The correlations that differ between the sexes appear in darker colors. (**H**) Interaction between NCI and mild cognitive impairment (MCI) and AD represented for the top 15 most significant interactions computed on the 100 most different pairs of subunits.

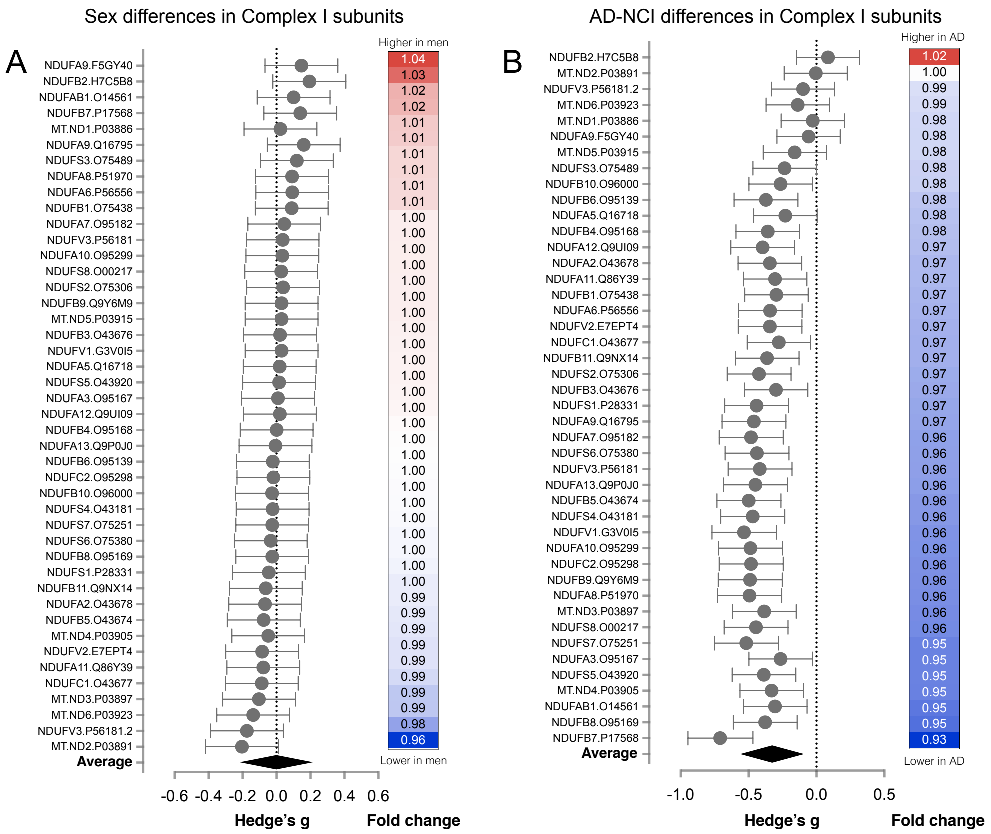


**Fig. S17. Difference in mitochondrial respiratory chain (RC) complex I proteins abundance according to sex and clinical diagnosis.** (**A**) Difference between men and women in complex I proteins abundance shown as (*left*) effect size and (*right*) fold change. (**B**) Difference between individuals with no cognitive impairment (NCI) and AD at death in RC complex I proteins abundance shown as (*left*) effect size and (*right*) fold change. The pooled effect size for AD vs NCI is -0.34 (95%IC, -0.58; -0.11). Detailed results are shown in supplementary table S4.
